# Repeated genomic signatures of adaptation to urbanisation in a songbird across Europe

**DOI:** 10.1101/2020.05.05.078568

**Authors:** Pablo Salmón, Arne Jacobs, Dag Ahrén, Clotilde Biard, Niels J. Dingemanse, Davide M. Dominoni, Barbara Helm, Max Lundberg, Juan Carlos Senar, Philipp Sprau, Marcel E. Visser, Caroline Isaksson

**Author notes:** Correspondence to Caroline Isaksson. Both authors contributed equally (Pablo Salmón, Arne Jacobs).

## Abstract

Urbanisation is currently increasing worldwide, and there is now ample evidence of phenotypic changes in wild organisms in response to this novel environment, but the extent to which this adaptation is due to genetic changes is poorly understood. Current evidence for evolution is based on localised studies, and thus lacking replicability. Here, we genotyped great tits (*Parus major*) from nine cities across Europe, each paired with a rural site, and provide evidence of repeated polygenic responses to urban habitats. In addition, we show that selective sweeps occurred in response to urbanisation within the same genes across multiple cities. These genetic responses were mostly associated with genes related to neural function and development, demonstrating that genetic adaptation to urbanisation occurred around the same pathways in wildlife populations across a large geographical scale.

## Main

Urban development is rapidly expanding across the globe, and although urbanisation is regarded a major threat for wildlife (Hendry, Gotanda, and Svensson 2017), its potential role as an evolutionary driver of adaptation has not been explored until recently (Johnson and Munshi-South 2017; J. C. Mueller et al. 2013; Harris and Munshi-South 2017; Rivkin et al. 2019). Some species have phenotypically adapted to the many urban challenges, such as higher levels of noise, artificial light at night, air pollution, altered food sources and habitat fragmentation (Alberti et al. 2017). Indeed, there is now evidence that some of these adaptations may have a genetic basis (Campbell-Staton et al. 2020), in line with the finding that such micro-evolutionary adaptations can occur within short timescales, particularly in response to human activities (Bosse et al. 2017; Hendry, Farrugia, and Kinnison 2008). However, the short evolutionary timescale, the dependence of evolution on local factors, and the polygenic nature of many phenotypic traits, make detecting evolutionary signals difficult (Hendry, Farrugia, and Kinnison 2008; Pritchard, Pickrell, and Coop 2010). Thus, we still lack important knowledge on the signals of adaptation, and thereby of the actual magnitude of the evolutionary change induced by urbanisation on wildlife populations.

The majority of available studies on the genetic bases of urban adaptation have either focused on a limited number of markers and genes (J. C. Mueller et al. 2013) or focused on a narrow geographical scale (Harris and Munshi-South 2017; Campbell-Staton et al. 2020; Perrier et al. 2018). As a result, an important gap remains in the understanding of the prevalence of convergent evolution among cities (Rivkin et al. 2019), limiting the inferences that can be made on the genomic response to urbanisation. A robust approach to address this gap is the use of paired urban and rural sites at a large spatial scale in combination with high throughput genomic tools. Such a strong replicated approach can help to simultaneously detect subtle allele frequency shifts and identify genomic regions repeatedly involved in parallel evolutionary adaptation or under divergent selection across distant urban habitats. This approach provides a powerful framework to test the repeatability of the genomic adaptation to urbanisation.

In this study, we present a multiple location analysis of the evolutionary response to urbanisation, using the great tit (*Parus major*). This widely-distributed passerine bird is a model species in urban, evolutionary and ecological research (e.g. Boyce and Perrins 1987; Charmantier et al. 2008; 2017; Pettifor, Perrins, and McCleery 1988; Bouwhuis et al. 2009; Krebs 1971; Salmón et al. 2017; Sprau, Mouchet, and Dingemanse 2017; Senar et al. 2017; Isaksson et al. 2009), with demonstrated phenotypic changes in response to urban environments in several populations (Charmantier et al. 2017; Sprau, Mouchet, and Dingemanse 2017; Senar et al. 2017; Caizergues, Grégoire, and Charmantier 2018). Additionally, genomic resources are well developed for this species (Kim et al. 2018) and it is known that across its European range, the species presents low genetic differentiation (Perrier et al. 2018; Laine et al. 2016; Lemoine et al. 2016; Spurgin et al. 2019). In order to examine and test the repeatability in the genomic responses to urbanisation in a broad geographical scale we analysed nine paired urban and rural great tit populations across Europe (Figs. 1a and 1b; Table S1). All urban sites used in the study were located in built-up areas or parks within the city boundaries, while rural sites were always natural or semi-natural forests containing only a few isolated houses. Additionally, we quantified the relative degree of urbanisation for each site (urbanisation score: PC_urb_, from principal component analysis, PCA; see Methods and Materials, Table S1; Fig. 1b). We combine two complementary approaches, the detection in urban habitats of parallel allele frequency shifts across many loci and the search for independent and repeated selective sweeps (i.e. a strong increase in haplotype frequency at one or few loci). Furthermore, we use functional enrichment analyses of genes putatively under selection to infer which particular phenotypes, known and unknown, are associated with genes under selection in urban great tits, providing an outline of genes to target in future ecological and functional studies.

**Fig. 1.**
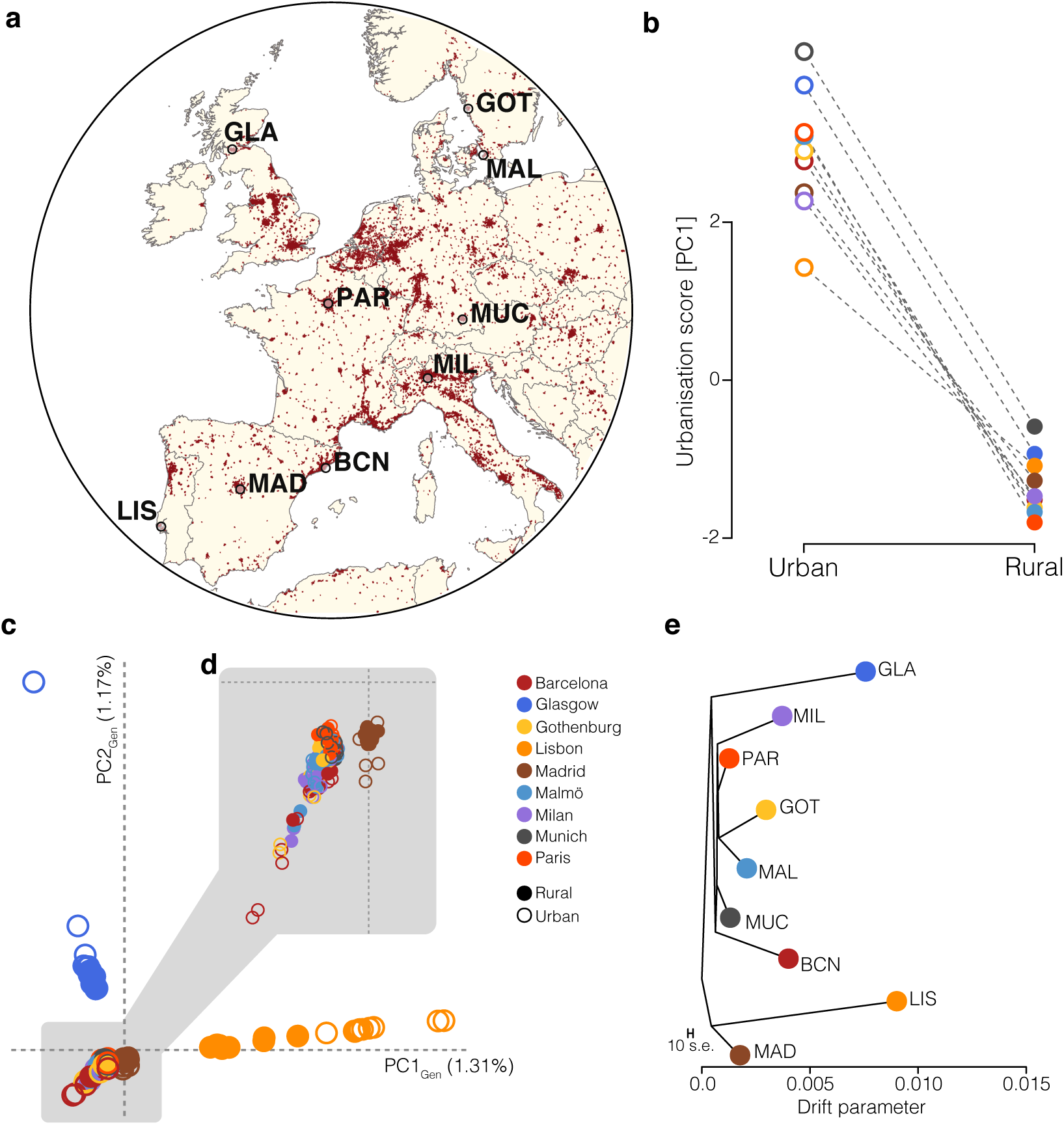
Urbanisation and population structure. **a**, Map of Europe, showing the targeted cities where the sampling of great tits (*Parus major*) was carried out. Red area indicates main dense urban areas. Shapefiles obtained from Natural Earth (https://www.naturalearthdata.com). **b**, Urbanisation scores (principal component, PC_urb_) for all nine urban-rural pairings. **c, d**, Principal component (PC_Gen_) plot showing the main axes of population structure for European great tits; dots represent individual birds. **d**, Zoomed view into the population cluster highlighted by the grey-shaded area. **e**, Maximum-likelihood tree showing the relationship between tits across sampling localities. BCN: Barcelona; GLA: Glasgow; GOT: Gothenburg; LIS: Lisbon; MAD: Madrid; MAL: Malmö; MIL: Milan; MUC: Munich; PAR: Paris.

## Results and Discussion

### Genetic diversity and population structure across European urban and rural populations

A total of 192 individuals were genotyped at 526,528 filtered SNPs, with 10-16 individuals per site (Table S1). The genetic diversity between each urban-rural pairing was relatively low, with genome-wide differentiation (F_ST_) ranging between 0.004 to 0.050 (Fig. S1c; Table S1). This is in line with previous studies of the species (Perrier et al. 2018; Laine et al. 2016; Lemoine et al. 2016; Spurgin et al. 2019). In addition, the levels of heterozygosity were similar between urban and rural populations, although slightly lower in some of the urban populations (see Table S1 for details; Wilcoxon test: W=30, P = 0.377). This indicates that the colonisation of urban environments is likely not a result of strong bottlenecks and its associated loss of genetic diversity. Despite the low population structure (Laine et al. 2016), two populations at the edge of the species distribution range (Lisbon and Glasgow) separated from all other populations along PC1_Gen_ and PC2_Gen_ (Fig. 1c; Figs. S1a and S2b). This pattern is possibly a consequence of the slightly reduced heterozygosity in these two populations (Figs. 1d and S3; Table S1). However, overall the population structure analyses suggest that urban-rural population pairs from the different localities have colonised each urban habitat independently, as urban-rural pairs split separately along PC_Gen_ (Fig. S2a), though still certain number of the urban populations cluster together.

### Identification of key SNPs associated with urbanisation

Despite the overall lack of genome-wide differentiation between urban-rural pairings, the selective pressures associated with urbanisation might have led to detectable genomic signatures of local adaptation. We used two complementary approaches, *LFMM* (Latent Factor Mixed Models) and an additional Bayesian approach (using *BayPass*), to narrow down genomic regions with consistent and strong allele frequency shifts associated with urbanisation. Testing for genotype-environment associations with *LFMM* (using PC_urb_ as a continuous habitat descriptor, Fig. 1b) revealed 2,758 SNPs associated with urbanisation (0.52% of the full SNP dataset, false-discovery rate (FDR) < 1%; Fig. 2a). These SNPs were widely distributed across the genome and did not cluster in specific regions. Larger chromosomes contained more urbanisation-associated SNPs (R^2^ = 0.97; Fig. 2c), highlighting the polygenic nature of urban adaptation. A PCA based on these SNPs clearly separated urban and rural populations along PC1_LFMM_ (Proportion of variance explained (PVE) by PC1 = 1.98%; Fig. S4a), showing highly parallel allele frequency changes in those loci across European cities. *BayPass* identified 70 urbanisation-associated SNPs (Bayes factor ≥ 20; Fig. 2b), of which 34 were shared with the *LFMM* analysis. These shared SNPs, which we term “*core urbanisation SNPs”* (Fig. 2d; Table S2), are likely involved in the local adaptation of great tits to urban habitats, and indeed, they strongly discriminated urban and rural individuals across Europe (PVE by PC1_GWAS_= 11.7%; Figs. 2e and S4b).

**Fig. 2.**
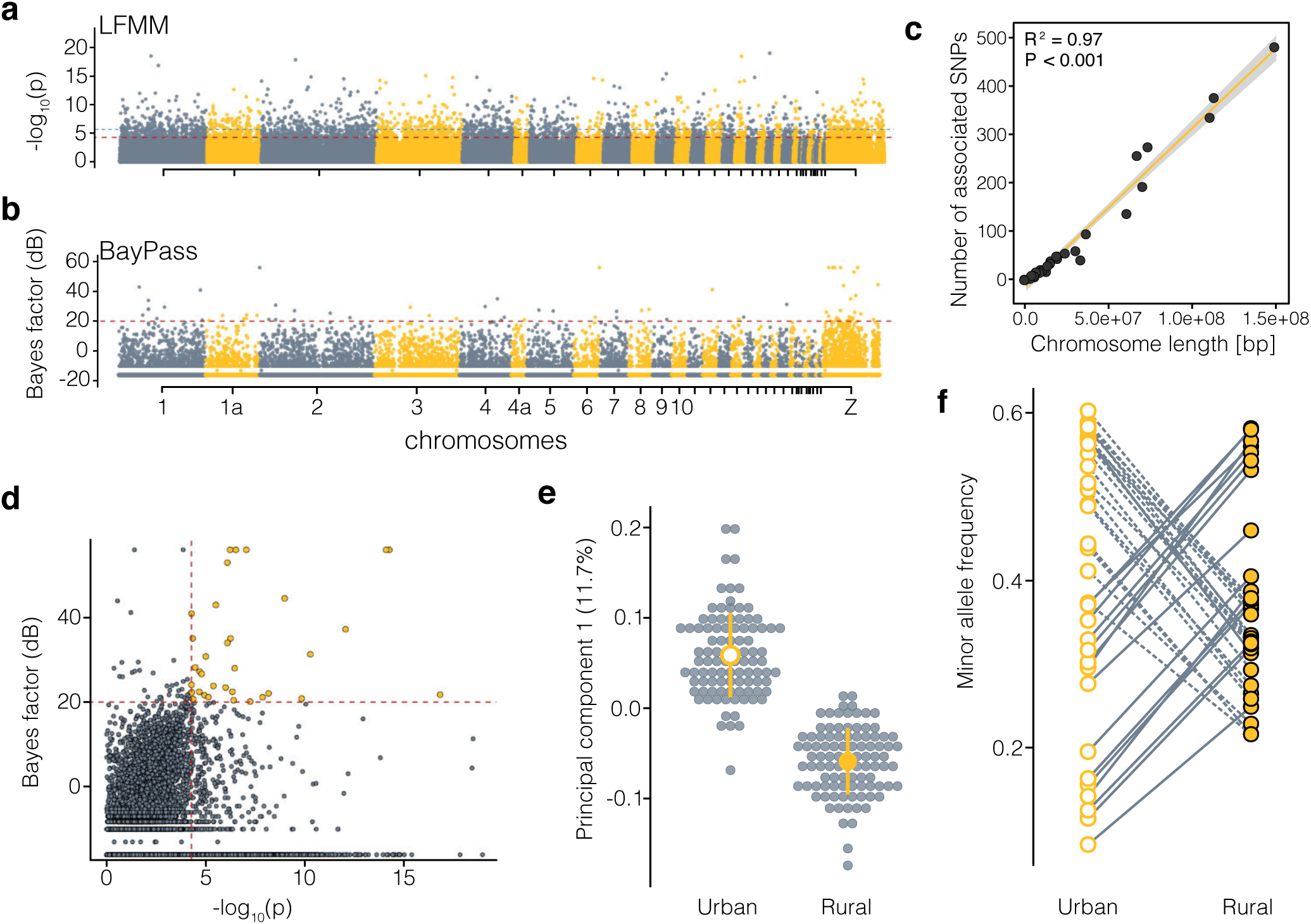
Genome-wide association with urbanisation. **a** and **b**, Manhattan plots showing signals of genome-wide association with urbanisation across all populations for the **a**, *LFMM* and **b**, *BayPass* analyses, respectively (see in methods “*Environment-associated SNPs*”). The red dotted line and the blue line in the *LFMM* Manhattan-plot show the 0.1 and 1% FDR significance thresholds. The red dotted line in the *BayPass* plot shows the significance threshold for a Bayes factor of 20 deciban (dB); alternating colors denote chromosomes. **c**, Correlation between the number of SNPs associated with urbanisation *per* chromosome in the *LFMM* analyses and the respective chromosome length. The strong correlation indicates a polygenic basis of urban adaptation. **d**, Correlation between association signals in *LFMM* and *BayPass*. The red dotted lines show the respective significance thresholds (*BayPass*: 20db; *LFMM*: 1% FDR). SNPs associated with urbanisation in both analyses are highlighted in yellow (“*core urbanisation SNPs*”, N=34). **e**, Main axis of variation in a PCA_GWAS_ based on “*core urbanisation SNPs*”. Grey circles show individuals and yellow dots and lines, show the mean ± s.d. for urban and rural populations. **f**, Reaction norm plot showing the difference in the mean minor allele frequency for “*core urbanisation SNPs*” between all urban and rural populations combined. See Fig. S5 for allele frequency trajectories by locality and SNP.

The importance of habitat (i.e. urban *versus* rural) was further underpinned by a univariate linear model. Using the first principal component axis of urbanisation-associated SNPs (PC1_GWAS_), habitat explained 73% of the total associated variance in allele-frequency divergence (η^2^_PC1 GWAS, Habitat_ = 0.73, P < 0.001). In comparison, both the effect of locality, which corresponds to the distinct evolutionary history of local populations (η^2^_PC1 GWAS, Locality_ = 0.20, P < 0.001), and the interaction of locality and habitat, which describes differences in the direction of allele-frequency change across cities (η^2^_PC1 GWAS, Habitat × Locality_ = 0.13, P = 0.001), explained much lower proportions of the variance than the habitat term on itself. Interestingly, the trajectories of minor allele frequency changes for the individual “*core urbanisation SNPs”* were highly repeatable across populations (Fig. S5). Thus, the directionality and/or magnitude in allele frequency of the 34 identified SNPs showed a highly parallel pattern across all urban populations (Fig. 2f), suggesting that in this species, local adaptation to urban habitats has occurred through repeated shifts in allele frequency of the same loci. This finding implies an important role of standing genetic variation in urban adaptation of great tits, as putatively adaptive alleles were shared across large parts of Europe (Spurgin et al. 2019).

### Signatures of selection in urban populations

Next, we performed a genome-wide scan of differentiation and selective sweep analyses to identify putative signatures of divergent selection between urban and rural populations. We first estimated genetic differentiation (F_ST_) in 200 kb sliding windows and 50 kb steps across the genome. While the genome-wide level of differentiation was low in all populations, we detected highly variable landscapes of genetic differentiation between adjacent urban and rural populations, with multiple genomic peaks of increased genetic differentiation (F_ST_ > 99^th^ percentile) spread across the genome (Fig. S6). 85 genomic windows were significantly differentiated in at least three urban-rural pairs.

Genetic differentiation can be driven by a myriad of processes, including background selection (selection against deleterious variants) or divergent selection (e.g. associated with local adaptation) in urban and/or rural populations (Burri 2017). To narrow down these processes, we determined whether the outlier windows were putatively under selection in urban populations by estimating population branch statistics (*PBS*) for each urban population in the same genome-wide windows. In this study, positive *PBS* values show an extended genetic distance of the urban population compared to the adjacent rural population and outgroup in a specific genomic window, indicative of positive selection in the urban population at that site (Yi et al. 2010). We used the rural population from Lisbon as the outgroup for estimating *PBS*, except for Lisbon, for which we used the rural population from Glasgow. We tested the effect of outgroup on *PBS* values but found that these were generally significantly correlated for each city (average spearman’s r = 0.525 ± 0.087 s.d., Fig S7 and S8). Between 25% and 50% of differentiated genomic regions (F_ST_) showed signs of selection based on *PBS* (top 1% of empirical *PBS* distribution) in urban populations (Fig. 3). Taking into consideration the nature of the urbanisation phenomenon, we might expect that the distinct genomic backgrounds would give rise to independent selective sweeps across localities, i.e. the result of locality-specific selection pressures. Yet, the shared genetic variation in this species across Europe(Spurgin et al. 2019) might lead to the re-use of shared adaptive variants across geographically distant populations. *PBS* values were highly variable across locations and showed generally low correlations among each other (|r| < 0.28) suggesting that putative selective sweeps are geographically limited, i.e. locality-specific. Nonetheless, 75 genomic windows with elevated *PBS* values were shared across more than three urban populations (maximum of five localities, Fig 3). At the gene-level, we detected multiple genes in or close to windows putatively under selection (within ≤ 100kb) across urban populations, of which five genes showed signs of selection in five out of nine urban populations (Table S3).

**Fig. 3.**
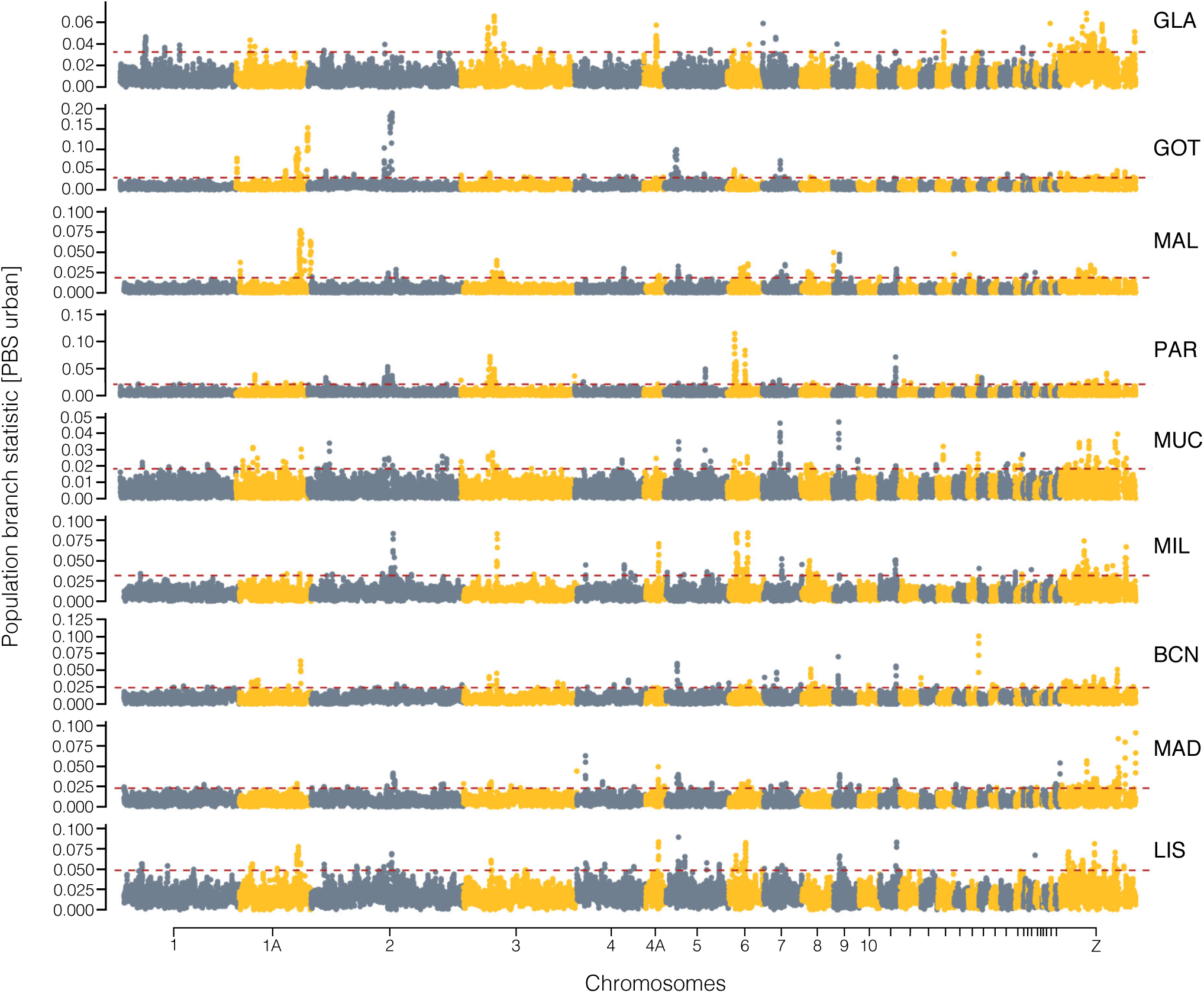
Shared signals of selection across European urban great tit populations. Population branch statistics (*PBS*) across the genome for each population in 200 kb sliding windows with 50 kb steps. Windows above the 99^th^ percentile of the empirical *PBS* distribution (red dashed line) were selected as putatively under selection in urban populations. Only positive values were plotted here for visualisation purposes. Full distributions can be found in Fig. S7 and S8. BCN: Barcelona; GLA: Glasgow; GOT: Gothenburg; LIS: Lisbon; MAD: Madrid; MAL: Malmö; MIL: Milan; MUC: Munich; PAR: Paris.

To complement the above analyses and to strengthen the inference of urban selective sweeps under divergent selection, we also used haplotype-based statistics, which are more robust to the effect of low recombination and linked selection (Tang, Thornton, and Stoneking 2007). We compared haplotype-homozygosity between each urban and its respective rural population (*Rsb score*) but only focused on absolute values as the ancestral states of haplotypes were not known (Meier et al. 2018). Weak Spearman pairwise-correlations of the haplotype-based *Rsb* scores for each SNP (Fig. S9 and S10) across localities indicate that selective sweeps are mostly locality-specific, as suggested by the weak correlations of *PBS*.

Nonetheless, we identified several genes with recurrent signals of selection across a broad geographical scale (> 3 urban populations). The *Rsb* statistics revealed signatures of selective sweeps in or close to 568 genes in more than three localities (4-8 urban populations), many of which were also detected using the *PBS* statistic (“*candidate genes under selection”*; Fig. 3d; Table S3). Comparing genes associated with signatures of selection by *PBS* and *Rsb* statistics, we identified 79 genes that showed signs of selection in at least three populations for both summary statistic (Table S3). Of these, 4 genes were putatively under selection in more than half of all urban populations (> 5 populations, maximum of 6) based on *PBS* scores, and also showed signs of selection in 5 to 8 populations (median = 5 ± 1.69 s.d.) based on *Rsb* scores (Table S3). The recurrent signals of selection on these genes suggest the existence of genetic parallelism in the underlying response to urbanisation at the gene level and indicate that these genes might potentially be strong candidates in the adaptation to urban habitats.

### Evolutionary drivers of genetic divergence and signatures f selection

To test if signatures of selection were caused by divergent selection, or alternatively by background selection in low recombination regions (Burri 2017), we evaluated the correlation of signatures of selection with patterns of linkage disequilibrium (LD). First, we tested the correlation between genetic diversity (LD measured as r^2^ and summarized in 200 kb sliding windows with 50 kb steps per population and summarised as PC1 across all urban populations) with measures of selection (*PBS* and *Rsb* summarised as PC1 across populations). Overall, the correlation between genetic differentiation (*PBS-PC1*) and genetic diversity in urban populations (*LD-PC1*) was relatively weak but significant (R^2^ = -0.002; P = 9.7 × 10^−11^), with low diversity regions (high LD) showing weaker signatures of selection, i.e. low *PBS* scores (Fig. 4a-c). This suggests that broad signatures of selection in urban habitats, positive *PBS-PC1*, are likely not caused by background selection in low recombination regions but rather divergent selection (Burri et al. 2015). We detected similar correlation patterns in most populations when we assessed the correlation of urban genetic diversity (*LD-PC1*) with *PBS* scores by locality, i.e. each independent city, rather than across all populations together (Fig. S11). However, in some localities, such as in Malmö (MAL) and Gothenburg (GOT), we detected a positive correlation between selection and genetic diversity (*PBS* ∼ *LD-PC1*), suggesting the existence of spatially varying impacts of background selection on the genomic landscape of differentiation in great tits (Fig. S11). Similarly, the correlation between haplotype-homozygosity (*Rsb-PC1*) and genetic diversity (*LD-PC1*) was very weak but significant (all chromosomes: R^2^ = 0.0115; P < 2.2 × 10^−16^, without Z chromosome: R^2^ = 0.0021; P = 6.9 × 10^−10^), with outlier windows not being strongly associated with low diversity regions (high LD) (Fig. 4d). This is expected, as the comparative nature of the haplotype-based *Rsb* score accounts for overall reduced diversity in shared low-recombination regions. Nonetheless, detailed recombination rate analyses and whole-genome data will be needed in the future to thoroughly understand the effect of recombination, linked selection and divergent selection on the landscape of divergence in European great tits.

**Fig. 4.**
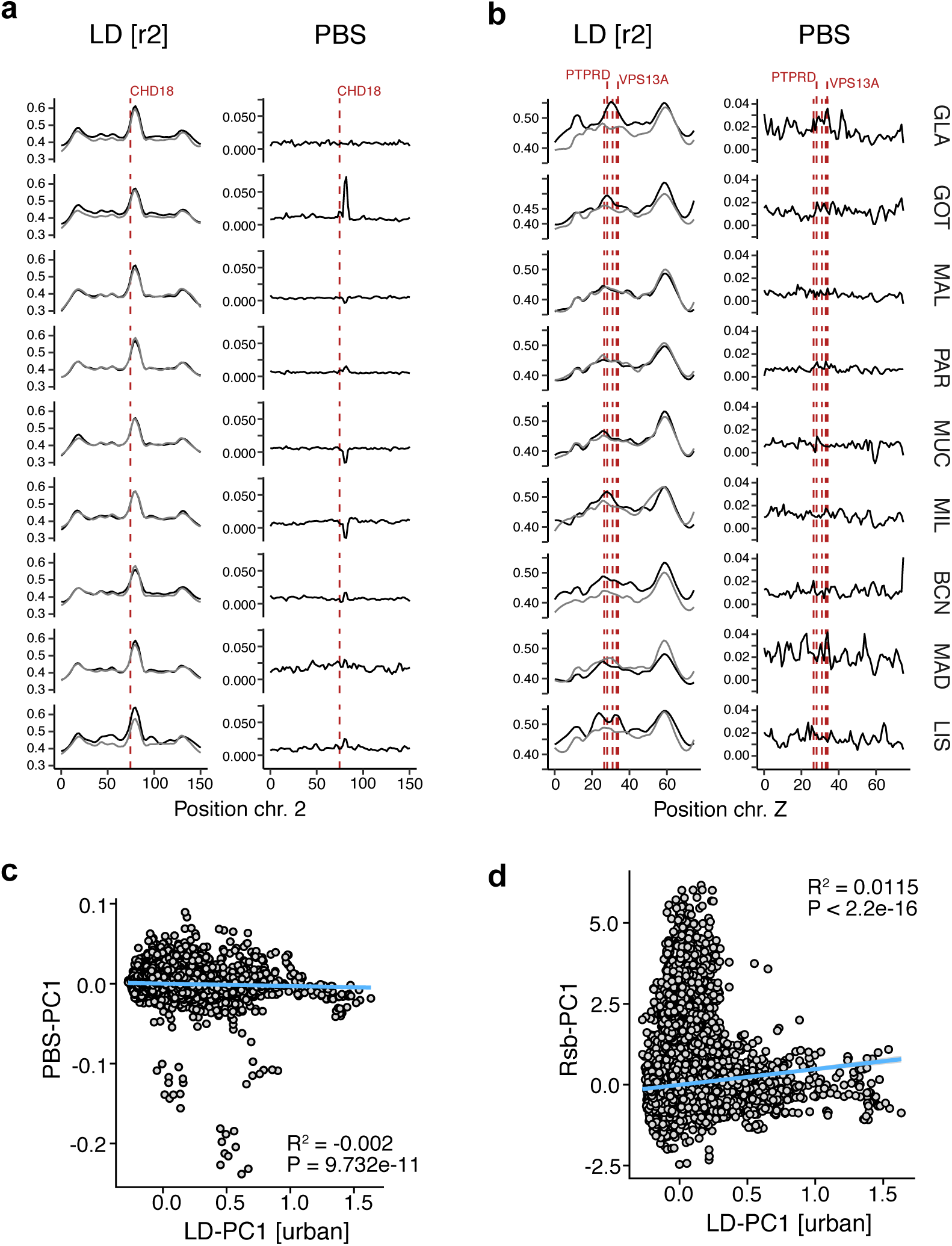
Signatures of selection and LD. **a, b** Patterns of LD and *PBS* around candidate genes associated with urbanisation on chromosome 2 and the Z chromosome. The positions of candidate genes are highlighted by red dashed lines. Patterns of LD for urban populations are shown in black and those for rural populations in grey. **c**, Correlation between *PBS-PC1* and *LD-PC1* (for urban populations) based on 200 kb sliding windows. **d**, Correlation between *Rsb-PC1* and *LD-PC1* (urban) based on the same 200 kb sliding windows. Note that windows with high *Rsb* values but low values of *LD-PC1* are located on the Z chromosome. See text for details. BCN: Barcelona; GLA: Glasgow; GOT: Gothenburg; LIS: Lisbon; MAD: Madrid; MAL: Malmö; MIL: Milan; MUC: Munich; PAR: Paris.

Furthermore, we assessed patterns of linkage disequilibrium around individual “*candidate genes under selection”* by locality (Figs. 4a and b). Low recombination regions or shared selective sweeps in urban and rural individuals would result in reduced genetic diversity in all populations (i.e. both urban and rural), as the recombination landscape in songbirds is generally highly conserved (Burri et al. 2015; Delmore et al. 2018). Yet, LD patterns around candidate genes differed strongly between chromosomes, suggesting effects of different evolutionary drivers (Fig. 4a and b), with most candidate regions not being located inside conserved low diversity regions (strongly correlated LD peaks) that might be indicative of a low recombination region. However, we found that one shared candidate gene (*CDH18*) that showed signatures of selection in five populations based on *Rsb* and *PBS*, was located inside a highly conserved LD peak on chromosome 2 (Fig. 4a, Table S3). This location potentially represents the centromere, and thus we cannot fully exclude the impact of background selection on the repeated signature of selection. However, the fact that *CDH18* only seems to show signatures of selection in urban populations but not rural populations suggests a role of divergent selection and adaptation to urban habitats.

Furthermore, LD was increased chromosome-wide in some urban populations compared to rural populations (e.g. chromosome Z, Fig. 4b), potentially suggesting an overall reduced genetic diversity and increased drift on those chromosomes. Nonetheless, local LD-peaks were still present around the “*candidate genes under selection”* in urban populations compared to rural populations (e.g. genes *PTPRD* and *VPS13A* in chromosome Z in Glasgow and Barcelona; Fig 4b, Fig. S12). The same genes also showed signs of selective sweeps in urban populations from localities without differences in baseline LD (e.g. chromosome Z in Milan; Figs. 4b), further supporting a scenario of divergent selection rather than increased drift due to locally reduced effective population size. Indeed, in birds, patterns of genetic differentiation at early stages of divergence, as presumably during colonisation of urban areas, have been found to be driven by selection rather than recombination (Burri 2017; Delmore et al. 2018). Thus, it is most likely that divergent selection rather than linked selection or drift explains the observed recurrent signals of selection in shared genes.

Interestingly, many of the SNPs putatively under selection were located on the Z-chromosome, which in birds is the sex-chromosome in the homogametic sex (males) (Figs. 3 and 4b). The strong urban association and selection signatures on this chromosome are in line with the “*Fast-Z*” evolution (Dean et al. 2015), which could be, at least partially, explained by a reduced recombination and effective population size on this particular chromosome. However, because many of these variants show similar allele frequency shifts between habitats across localities (Fig. 2f) and show signatures of selection in urban but not rural populations, recombination differences on their own cannot be the main driving factor of divergence in this particular chromosome.

### Convergent evolution of gene functions in response to urbanisation

While the majority of genes showing signs of selection were unique to one or two cities, supporting the scenario of independent selective events, a significant proportion of genes were putatively under selection in multiple cities. Indeed, permutation analyses indicated that one would not expect, by chance, the same gene under selection in three or more cities (χ^2^_1_ =77.947, P < 2.2 × 10^−16^). However, 11 individual genes showed signs of selective sweeps in more than half of the studied urban populations (five to six) based on *PBS*, and also in more than three populations (three to eight) based on the *Rsb* statistic (Fig 5a, Table S3). Additionally, 107 genes were putatively under selection in three or four urban populations based on *PBS* and also showed signs of selection based on *Rsb* in more than three populations (three to seven) (Table S3). These results corroborate and reinforce evidence for repeatability at the genomic level in the response to urbanisation between distant urban populations. Furthermore, 14 of these 79 genes putatively under selection were also associated with urbanisation-associated SNPs (see *LFMM* analyses).

**Fig. 5.**
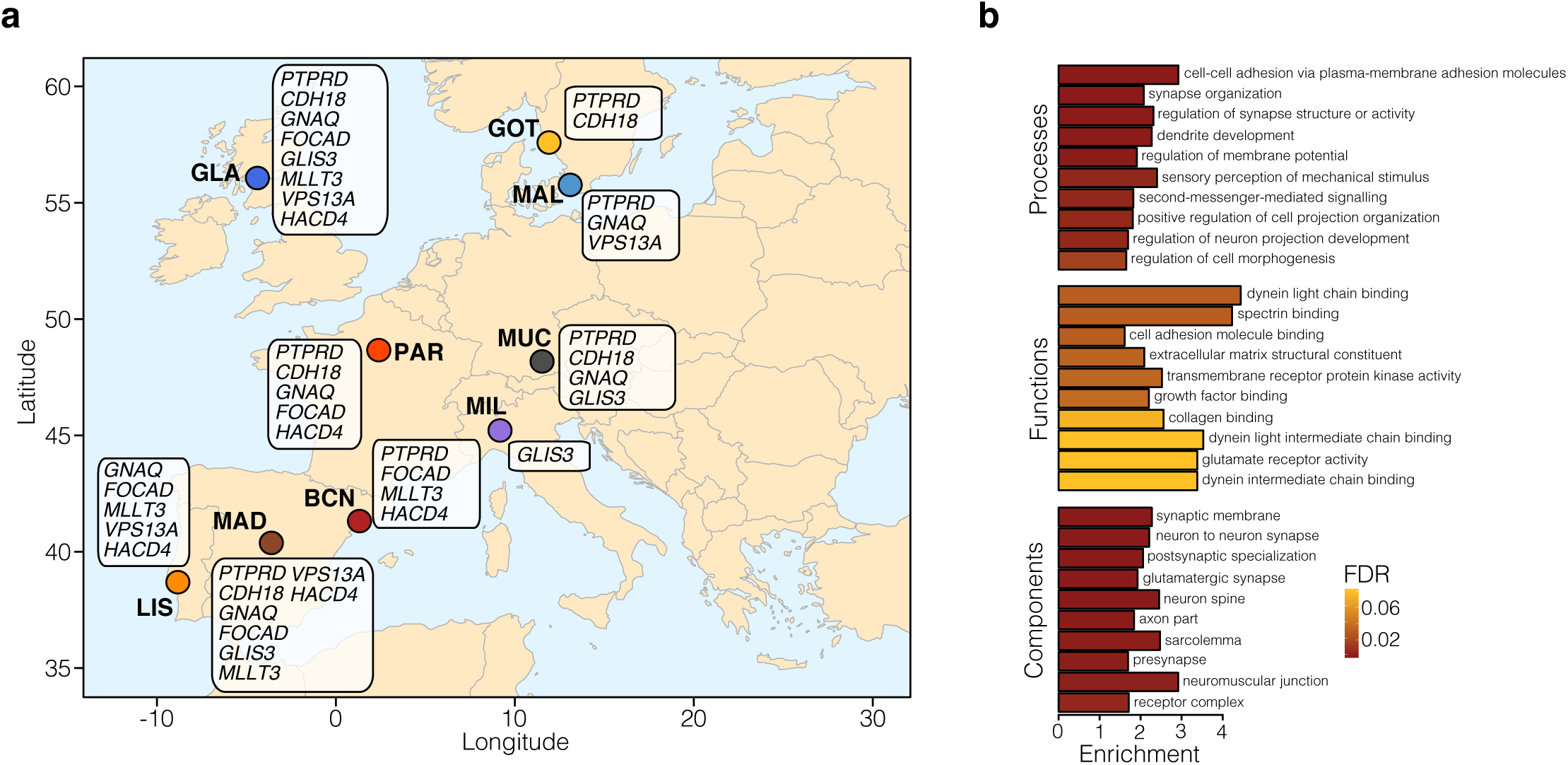
Candidate genes and biological pathways associated with urbanisation. **a**, Map showing the spatial distribution of some top shared candidate genes putatively under selection in urban populations across Europe. **b**, Enrichment of associated genes in significantly enriched gene ontology (GO) groups by GO category in the *LFMM* and *BayPass* analysis (see Table S4 for a detailed list). The colour of bars represents the false-discovery rate (FDR) from the gene ontology overrepresentation analyses. BCN: Barcelona; GLA: Glasgow; GOT: Gothenburg; LIS: Lisbon; MAD: Madrid; MAL: Malmö; MIL: Milan; MUC: Munich; PAR: Paris.

Many of the genes associated with urbanisation and under recurrent selective sweeps in urban populations have previously been linked with behavioural divergence, suggesting adaptive phenotypic shifts related to behaviour. For instance, the *PTPRD* gene (chr. Z), which showed signs of selection in seven urban populations and was associated with urbanisation (Figs. 5b, Fig S12), encodes for the receptor-type tyrosine-protein phosphatase delta, an enzyme suggested to be involved in neural development of the hippocampus (Uetani et al. 2000), a brain region linked to spatial memory, bird navigation and flight performance (Mehlhorn, Haastert, and Rehkämper 2010; Gazda et al. 2018). The *CDH18* gene (chr. 2) is part of a superfamily of membrane proteins involved in synaptic adhesion and was revealed as a candidate gene in phonological alterations in humans (Peter et al. 2016). Furthermore, *VPS13A* (chr. Z) gene variants in humans are linked to chorea-acanthocytosis (Ishida et al. 2009), a neurodegenerative disorder that affects movement and, this gene has recently shown to be associated with migratory behaviour in a songbird (Toews et al. 2019).

Thus, our selection analyses suggest that natural selection repeatedly acts on behavioural traits and sensory and cognitive performance, all previously shown to be among the most widespread differences between urban and rural wildlife populations (Sih and Del Giudice 2012; Sol, Lapiedra, and González-Lagos 2013). These findings were furthermore supported by the GO terms analysis of the 2,758 urbanisation-associated SNPs (*LFMM* analysis), linked to 984 genes (1,501 SNPs in genic regions). Accordingly, most of the GO terms were related to neural functioning and development (e.g., GO:0016358, FDR = 4.30 × 10^−6^), cell-adhesion (e.g., GO:0098742, FDR = 3.43 × 10^−6^) and sensory perception (e.g., GO:0050954, FDR = 3.42 × 10^−3^; Fig. 4, Table S4). These GO terms were mainly clustered into two interacting networks, one related to sensory recognition and the other to neural development and cell adhesion (Fig. S13). These findings reinforce the previous idea on the importance of cognitive and behavioural changes as key responses to urbanisation in birds and in particular in great tits. Indeed, song structure and escape or distress behaviour have been previously shown to differ between urban and rural great tit populations across Europe (Senar et al. 2017; Slabbekoorn and den Boer-Visser 2006; Møller and Ibáñez-Álamo 2012). Nonetheless, whether this is the result of a genetic response to selection or phenotypic plasticity is to a large extent still unknown. Only a few studies have previously explored the genetic patterns underlying urban adaptation in birds, finding evidence of divergence in behaviour-related genes at multiple European populations using either a candidate gene approach (J. C. Mueller et al. 2013) or a low-density SNP along transects within a city (Perrier et al. 2018). Furthermore, similar pathways showed divergence across three neighboring urban areas in a recently established urban-dwelling avian species in South America (Jakob C. Mueller et al. n.d.). Overall, our study supports these earlier observations regarding adaptive genetic changes in behavioural and neural development, suggesting that these processes play an important role in urban adaptation. Detailed functional genomic and phenotypic analyses are now needed to understand the role of these genes in the adaptive divergence of urban and rural great tits and other songbirds.

## Conclusions

Our study demonstrates genetic signals of repeated local adaptation to urban habitats in a common songbird across multiple European cities. We found that a combination of parallel polygenic allele frequency shifts and recurrent but independent selective sweeps are associated with adaptation to urban environments. Our results strongly suggest that a few genes with known neural developmental and behavioural functions experienced recurrent but independent selective sweeps only in the urban populations. This suggests a strong consistency in the processes associated with urbanisation, despite the fact that underlying haplotypes are not shared. Thus, our study exemplifies repeated evolutionary adaptation to urban environments on a continental scale and highlights behavioural and neurosensory adjustments as important phenotypic adaptations in urban habitats.

## Methods

### Sample collection and DNA extraction

During the years 2013-2015, 20 or more individual great tits were sampled at paired urban-rural sites from nine European cities (Figure 1a; Table S1). We sampled a total of 192 individuals (aged > 1 year old) with 10-16 individuals per site (Table S1). Sexes were balanced between pairs (urban-rural) in the dataset (GLMM; pair: χ^2^_1_ = 0.505, P= 0.477). Each of the paired sampling sites (urban or rural, hereinafter populations) was sampled within the same season. Barcelona and Munich were sampled during winter period, however, in both cases only known birds (recaptures) were included in the study, thus, all birds can be considered resident. All urban populations were located within the city boundaries, the areas are characterized with significant proportion of human-built structures such as houses and roads with managed parks as the only green space. Rural populations were chosen to contrast the urban locations, regarding degree of urbanisation and were always natural/semi-natural forests and contained only a few isolated houses. Each urban and rural population were separated with a distance above the mean adult and natal dispersal distance of this species (i.e. see Table S1) (Paradis et al. 1998).

Blood samples (approx. 25 µl) were obtained either from the jugular or brachial vein and stored at 4 °C in ethanol or SET buffer and subsequently frozen at -20 °C. In each case, procedures were identical for the paired rural and urban populations. DNA was extracted from approximately 5 µl samples of red blood cells in 195 µl of phosphate-buffered saline using Macherey-Nagel NucleoSpin Blood Kits (Bethlehem, PA, USA) and following the manufacturer’s instructions or manual salt extraction (ammonium acetate). The quantity and purity of the extracted genomic DNA was high and measured using a Nanodrop 2000 Spectrophotometer (Thermo Fisher Scientific) and Qubit 2.0 Fluorometer (Thermo Fisher Scientific).

### Urbanisation score

To quantify the degree of urbanisation at each site we used the *UrbanizationScore* image-analysis software, based on aerial images from *Google Maps* (Google Maps 2017) and following the methods previously described in different studies assessing the effect of urbanisation on wild bird populations (Seress et al. 2014). Briefly, each sampling site was represented by a 1 km x 1 km rectangular area around the capture locations. The content in each rectangle was evaluated dividing the image in 100 m x 100 m cells and considering three land-cover characteristics in each: proportion of buildings, vegetation (including cultivated fields) and paved surfaces. The different land-cover measures obtained per site were used in a principal component analysis to estimate an urbanisation score variable (PC_urb_) for each of the urban or rural populations per locality, see Supplementary Table S1. The PC_urb_ values were transformed to obtain negative values in the less urbanized and positive in the more urbanized sites. We used the average of the urbanisation estimates if birds were captured in more than one location within each site (>2 km apart, mean ± s.d.: 931.22 ± 1,005.26 m). All quantifications were done in triplicates by the same person (P.S.) and the estimates were highly repeatable, (intra class correlation coefficient, ICC = 0.993, 95% CI = 0.997-0.987, P < 0.001).

### SNP genotyping

All 192 individuals were successfully genotyped using a custom made Affymetrix© great tit 650K SNP chip at Edinburgh Genomics (Edinburgh, United Kingdom). SNP calling was done following the “*Best Practices Workflow*” in the software Axiom Analysis Suite 1.1.0.616 (Affymetrix©) and all the individuals passed the default quality control steps provided by the manufacturer (dish quality control values > 0.95) and previous studies using the same SNP chip (*6, 17*). A total of 544,610 SNPs were then exported to a variant-calling format (VCF) and Plink and furthered filter and assigned to chromosomes using the great tit reference genome (GCA_001522545.2 *Parus major* v1.1; NCBI Annotation Release 101). 155 SNPs were not found in the new assembly and 17,927 SNPs were not in chromosomic regions; thus, these SNPs were removed from further analysis leaving a total of 526,528 SNPs.

### Genetic diversity and Population Structure

We calculated the genome-wide genetic diversity as expected heterozygosity (H_e_) for each population using *Plink 1.9* (Purcell et al. 2007), and tested if genetic diversity significantly differed between urban-rural populations from the same location using t-tests in R and overall across urban-rural pairings using a Wilcoxon rank sum test in *R* (R Core Team 2018). Furthermore, we estimated pairwise F_ST_ between all population-pairs (urban-rural per locality) using *SNPRelate* (Zheng et al. 2012). Mean average F_ST_ was computed across all comparisons after setting negative values to zero (Zheng et al. 2012).

For analyses of population structure, we pruned the SNP dataset based on linkage disequilibrium (LD) in *Plink 1.9* using a variance inflation factor threshold of 2 (“*-indep 50 5 2*”), retaining 358,149 SNPs. Using this pruned and filtered dataset (314,350 SNPs), we performed a principal component analysis using *SNPRelate* (Zheng et al. 2012). Genetic ancestry analysis was done using the software package *Admixture v.1.3* (Pickrell and Pritchard 2012) with K ranging from 2 to 18 and ten-fold cross-validation.

Additionally, we inferred a population tree based on allele-frequency co-variances using *Treemix v.1.3* (Pickrell and Pritchard 2012), with blocks of 500 SNPs. In order to test for the potential of secondary gene flow across populations (cities) we fitted up to five migration edges and determined the best fitting tree based on the increase in maximum likelihood, variation explained, and on the significance of migration edges (Jacobs et al. 2020).

### Environment-associated SNPs

We used two different approaches to identify SNPs associated with the degree of urbanisation. The degree of urbanisation was coded based on the “*urbanisation score*” to maintain gradual differences between rural and urban environments across the studied populations (see above). First, we used a univariate latent-factor linear mixed model implemented in *LFMM* for examining allele frequency – environment associations (Frichot et al. 2013). Based on the number of ancestry clusters (K) inferred with *Admixture* and the distribution of explained variation in the PCA (Fig. S2b), we ran *LFMM* with two and four latent factors, respectively. Each model was run 5 times for 10,000 iterations with a 5,000-iteration burn-in. We calculated the median z-score for each locus across all 10 runs and selected SNPs with a false-discovery rate (FDR) below 1% to be associated with urbanisation. The results with two or four latent factors were highly concordant and the same candidate loci were recovered; thus, we only used the results obtained with four latent factors for further analyses.

Second, we analysed associations with urbanisation using the auxiliary covariate model implemented *in BayPass v.2.1*. (Gautier 2015). We estimated the allele-frequency – environment association for each SNP with the urbanisation score for each population accounting for population structure using a covariance matrix. We estimated the covariance matrix using the LD pruned SNP dataset in the core model using default parameters: 20 pilot runs of 1,000 iterations, a run length of 50,000 iterations, sampling every 25^th^ iteration, and a burn-in of 5,000 iterations. The resulting covariance matrix was used as input for five replicated runs of the auxiliary covariate model using the above settings. The strength of association is given in the test by estimated Bayes factor (measured in deciban; dB). We calculated the median Bayes Factor across all five replicated runs and considered all SNPs with a deciban unit (dB) > 20 as urbanisation associated. This is the strictest criterion and is considered as “decisive evidence” for the association (Gautier 2015).

### Patterns of genetic differentiation (F_ST_, PBS)

To identify genomic regions distinguishing adjacent urban and rural great tits, we estimated the genetic differentiation (Weir & Cockerham’s F_ST_ (Weir and Cockerham 1984)) for each urban and rural pair for each SNP using *vcftools*. We subsequently summarised and plotted F_ST_ values in 200 kb sliding windows with 50 kb steps using the *windowscanr* R-package. To identify genomic regions putatively under selection in urban but not rural populations, we also calculated the population branch statistic (*PBS*) for each urban pair. We used the rural LIS (Lisbon) population as the outgroup as it is the genetically most distinct rural population, except for the *PBS* estimation for the urban LIS population, for which we used the rural GLA (Glasgow) population as the outgroup. We used the following formula from Zhang et al. (Zhang et al. 2020): PBS_urb_ = (T_urb-rur_ + T_urb-out_ – T_rur-out_) / 2, where T = -log(1-F_ST_). T_urb-rur_ is derived from F_ST_ between adjacent rural and urban populations, T_urb-out_ from F_ST_ between the focal urban and outgroup population, and T_rur-out_ from F_ST_ between the rural and outgroup population. For visualisation purposes, only positive *PBS* values, those showing putative signs of selection in the focal urban population, were plotted along the genome.

### Haplotype-based selection analysis

To identify genomic regions showing signs of (incomplete) selective sweeps in urban-rural population pairs, we scanned the genome for regions of extended haplotype homozygosity (EHHS) in urban compared to rural populations. We firstly used *fastPHASE* (Scheet and Stephens 2006) to reconstruct haplotypes and impute missing data independently for each chromosome using the default parameters, except that each individual was classified by its population (“*-u*” option). We used 10 random starts of the EM algorithm (“*-T*” option) and 100 haplotypes (“*-H*” option). The *fastPHASE* output files were analysed using *rehh 2.0* (Gautier, Klassmann, and Vitalis 2017). In addition, we used “*rehh*” to calculate *Rsb* statistics per focal SNP. The *Rsb* score is the standardized ratio of integrated EHHS (iES, which is a site-specific extended haplotype homozygosity) between two populations, is calculated using the following formula in *rehh* (Tang, Thornton, and Stoneking 2007; Gautier, Klassmann, and Vitalis 2017):

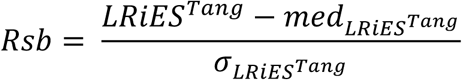

with LRiES^Tang^ representing the unstandardized log ratio of the iES^Tang^ _(urban)_ and iES^Tang^ _(rural)_ computed in the urban and rural populations (Gautier, Klassmann, and Vitalis 2017), and med_LRiES,Tang_ and ._LRiES,Tang_ representing the median and standard deviation of LRiES^Tang^, respectively. This statistic measures the extent of haplotype homozygosity between two populations and follows the rationale that if a SNP is under selection in one population compared to the other, the region around this locus will show an unusually high level of haplotype homozygosity compared to the neutral distribution. As the we don’t know the ancestral and derived state of each SNP, we focused on the absolute *Rsb* values. In accordance to Gautier and colleagues (Gautier, Klassmann, and Vitalis 2017), significant genomic regions were selected based on a threshold of |*Rsb*| ≥ 4. Because recombination rates can be assumed to be conserved between closely-related urban and rural populations, the cross-population comparative nature of the *Rsb* statistic provides an internal control that cancels out the effect of heterogeneous recombination across the genome (Tang, Thornton, and Stoneking 2007).

To determine the repeatability of selection across urban-centres, we implemented a resampling approach to assess the likelihood of genes showing signs of selection in two, three, four or more populations. We resampled with replacement n genes (n = number of genes with signatures of selection in each urban population) for each population from the list of all SNP-linked genes using the *resample* function in *R*, assessed the amount of overlap between populations (from two to 8 populations) and repeated the sampling 100,000 times for each comparison. We then calculated the mean and 95% confidence interval (CI) for each comparison and compared the number of observed shared candidate genes to the expected number of candidate genes. The expected number of genes showing signs of selection in three or more populations was zero, thus we focused on genes showing signs of selection (*Rsb, PBS*) in three or more populations. This highlights the fact that it is unlikely that genes repeatedly show signs of selection in three or more urban populations by chance alone.

### Parallelism analyses of urbanisation-associated SNPs

To determine the explanatory power of urbanisation associated SNPs and parallelism in allele frequency changes across populations, we performed a principal component analysis based on different candidate subsets, *i)* all *LFMM* candidate SNPs (PC_LFMM_ and *ii)* those overlapping between the *LFMM* and *BayPass* analyses (“*Core urbanisation SNPs*”, PC_GWAS_, see above), using *SNPrelate*. Furthermore, we ran a univariate linear model for each of the first three principal components (i.e. “PC _(PC1)_ ∼ Habitat + Locality + Habitat × Locality”) to quantitatively test the effect of parallel (significant “Habitat” effect, urban-rural) and non-parallel (significant “Habitat × Locality”) on allele-frequency changes across populations. We used the “*EtaSq*” function implemented in *BaylorEdPsych* (Beaujean and Beaujean, n.d.) to extract the effect sizes (partial η^2^) for the model terms in each linear model.

### Patterns of linkage disequilibrium (LD) across the genome

To estimate the impact of variation in recombination rate and linked selection in low-recombination regions on patterns of divergence (i.e. regions showing signs of selective sweeps), we estimated patterns of LD (r^2^) across the genome using *Plink 1.9* (Purcell et al. 2007). We focused on long-distance LD by calculating LD for pairs of SNPs within 2,000 to 200,000bp of each other for each population (each rural and urban population per locality). To reduce the computational load, we randomly sampled 5% of the dataset for each locality and plotted them for chromosomes containing candidate genes showing signs of selective sweeps in four or more cities (chromosome 2, chromosome Z). Furthermore, to investigate more large-scale patterns of correlation between selection and LD, we estimated a PCA based on LD scores in 200 kb sliding windows (50 kb steps) and used PC1 (*LD-PC1*) as a proxy for the distribution-wide LD pattern across the genome. We then estimated the PCA for PBS and estimated the correlation between *LD-PC1* and *PBS-PC1*. We also estimated the correlation between *LD-PC1* and *PBS* by locality to investigate local differences. We performed the same analyses for sliding window *Rsb* scores.

### Functional characterization of candidate SNP

We obtained the gene annotations for all candidate SNPs from the great tit reference genome annotation (GCA_001522545.2 Parus_major1.1; NCBI Annotation Release 101). We used all genes containing SNPs associated with urbanisation (*LFMM* and *BayPass)* (N = 1,501 SNPs within genes). To analyse the enrichment of functional classes, we identified overrepresented gene ontologies (biological processes, molecular functions and cellular components) using the *WebGestalt* software tool (Wang et al. 2017). The gene background was set using annotated great tit genes (Annotation release 101) containing SNPs from the SNP chip and with *H. sapiens* orthologues. *H. sapiens* genes were used as a reference set as human genes are better annotated with GO terms than those of any avian system (e.g. chicken) (Bosse et al. 2017; Laine et al. 2016). We focused on non-redundant gene ontology (GO) terms to account for correlations across the GO graph topology and GO terms as implemented in *WebGestalt* (Wang et al. 2017). A FDR < 0.05 was used as a threshold for significantly enriched GO terms. Furthermore, we searched the public record for functions of individual candidate genes. We also used *GOrilla* (Eden et al. 2009) to visualize the connections of GO terms associated with LFMM candidate genes and *Cytoscape v.3.6.1* (Shannon et al. 2003) to visualize the GO network and identify all enriched GO terms (biological processes, P < 0.001), including redundant terms.

Candidate genes associated with signatures of selection were those that were either overlapping with PBS outlier windows or within 100 kb up- or down-stream of a SNP with an absolute Rsb score above 4. Overlap between genomic regions/SNPs and genes was assessed using *bedtools*.

## Acknowledgements

We are truly thankful to J. Pérez-Tris, J. Ignacio Aguirre de Miguel, M. Morganti, ringers from La Herreria and Lago di Pusiano, V. Encarnação, A. Mouchet and Pardal family. We also thank V.N. Laine, A. Herrera-Dueñas and A.C. Mateman for their help during data extraction and formatting and laboratory work and M. Bosse for comments on an early version of the manuscript.

## Authors contribution

C.I. and P.S. conceived the study. P.S. collected and coordinated the sample collection with additions from C.B., N.D., D.M.D., B.H., J.C. and Ph.S. The data analysis was carried out by A.J. and P.S. with help from M.L., D.A. and M.E.V. C.I., A.J. and P.S. drafted the initial manuscript with input from all the other authors

## Supplementary Materials for

**Fig. S1.**
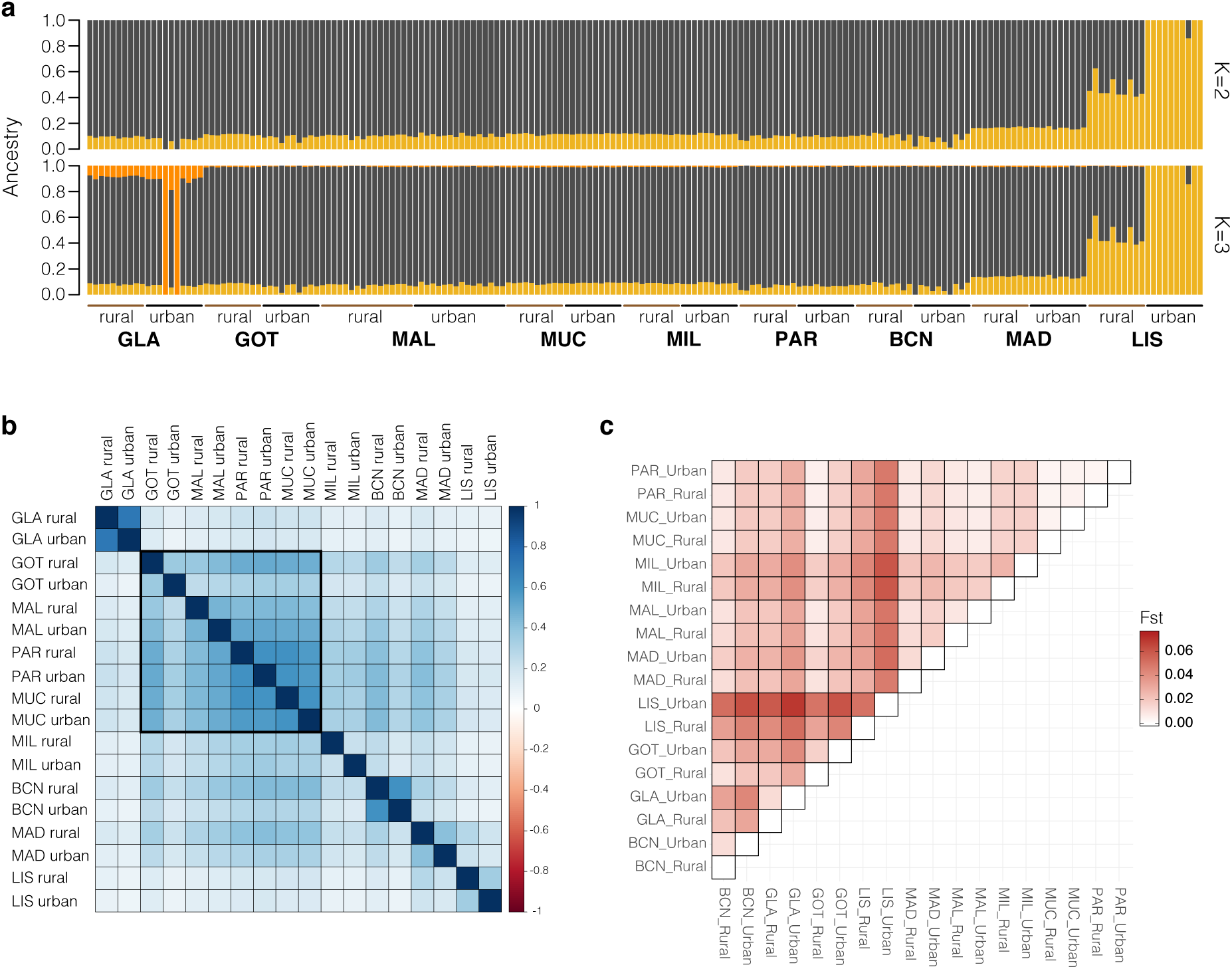
Genetic ancestry in urban and rural European great tits. **a**, Genetic ancestry inferred with *Admixture* for K=2 (CV-error = 0.615) and K=3 (CV-error = 0.618), showing two main genetic clusters across Europe. **b**, Genetic correlation matrix showing genome-wide genetic similarities among all studied populations inferred using *BayPass*. The colour gradient indicates the strength of the correlation. **c**, Heatmap of pairwise F_ST_ values between populations. BCN: Barcelona; GLA: Glasgow; GOT: Gothenburg; LIS: Lisbon; MAD: Madrid; MAL: Malmö; MIL: Milan; MUC: Munich; PAR: Paris.

**Fig. S2.**
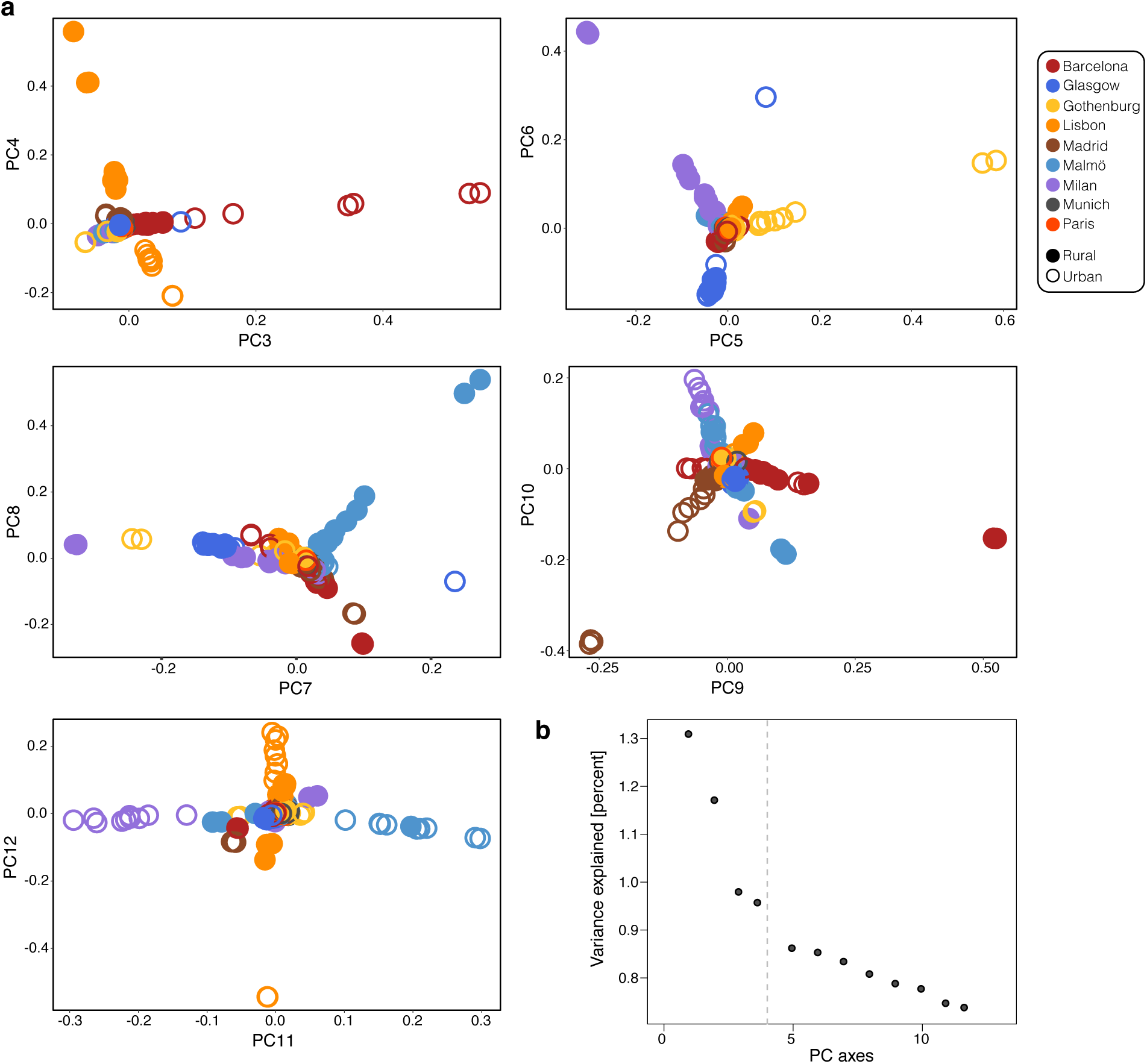
Population structure of European great tits. **a**, Principal component plots displaying the divergence of great tit populations along eigenvectors 3–12 (PC_Gen_). **b**, Percentage of total variance explained by the first 12 principal components. The dashed grey line indicates the number of eigenvectors used for population structure correction in *LFMM* based on the flattening of the distribution (K=2-4).

**Fig. S3.**
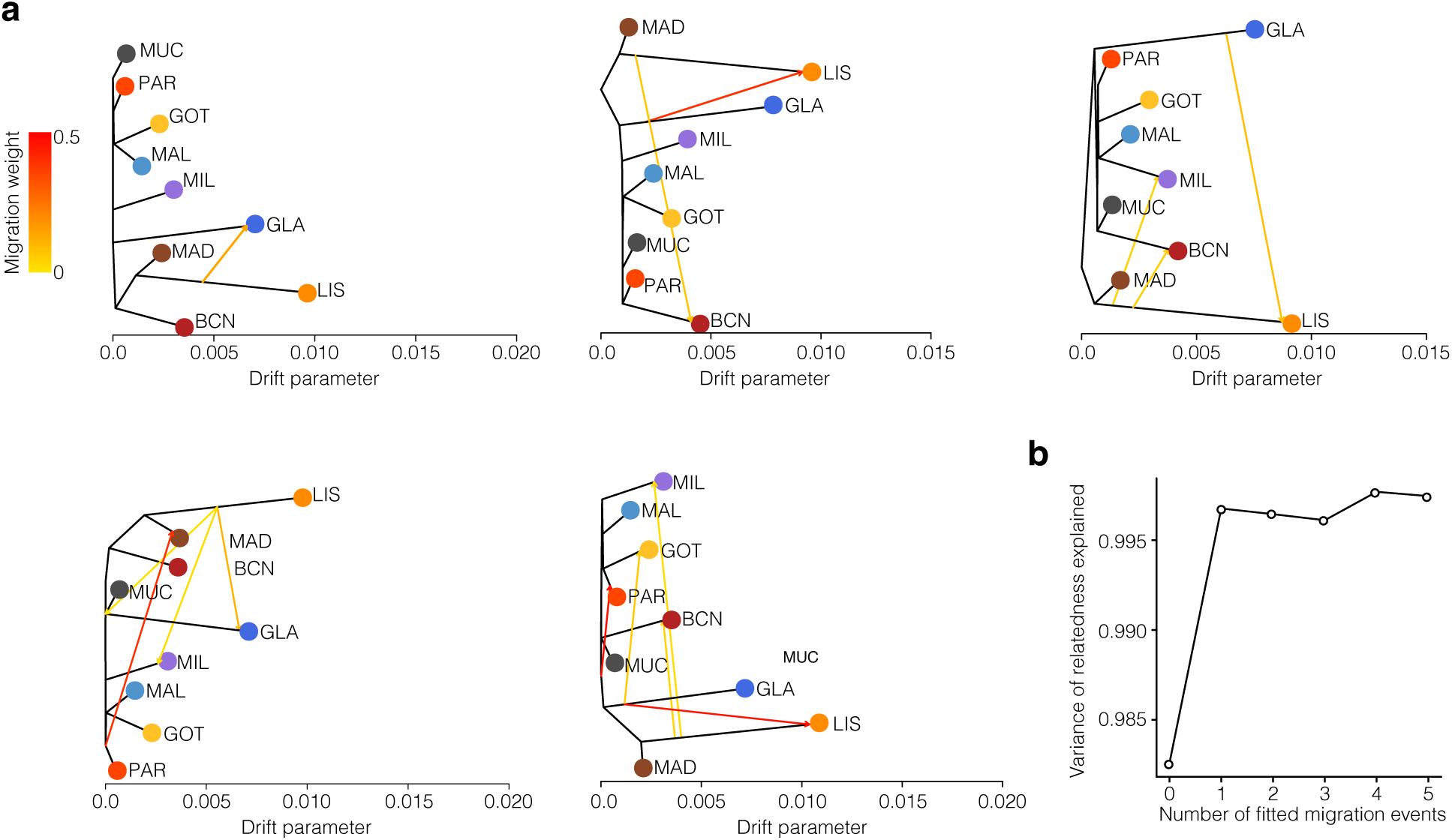
*Treemix* maximum-likelihood trees. **a**, Allele frequency based unrooted maximum-likelihood *Treemix* trees including one, two, three, four and five migration edges. The colour of the arrows shows the migration weight for gene flow events between branches. **b**, Proportion of variance explained by models with zero to five migration edges. Note the reduced rate of increase after adding one migration event.

**Fig. S4.**
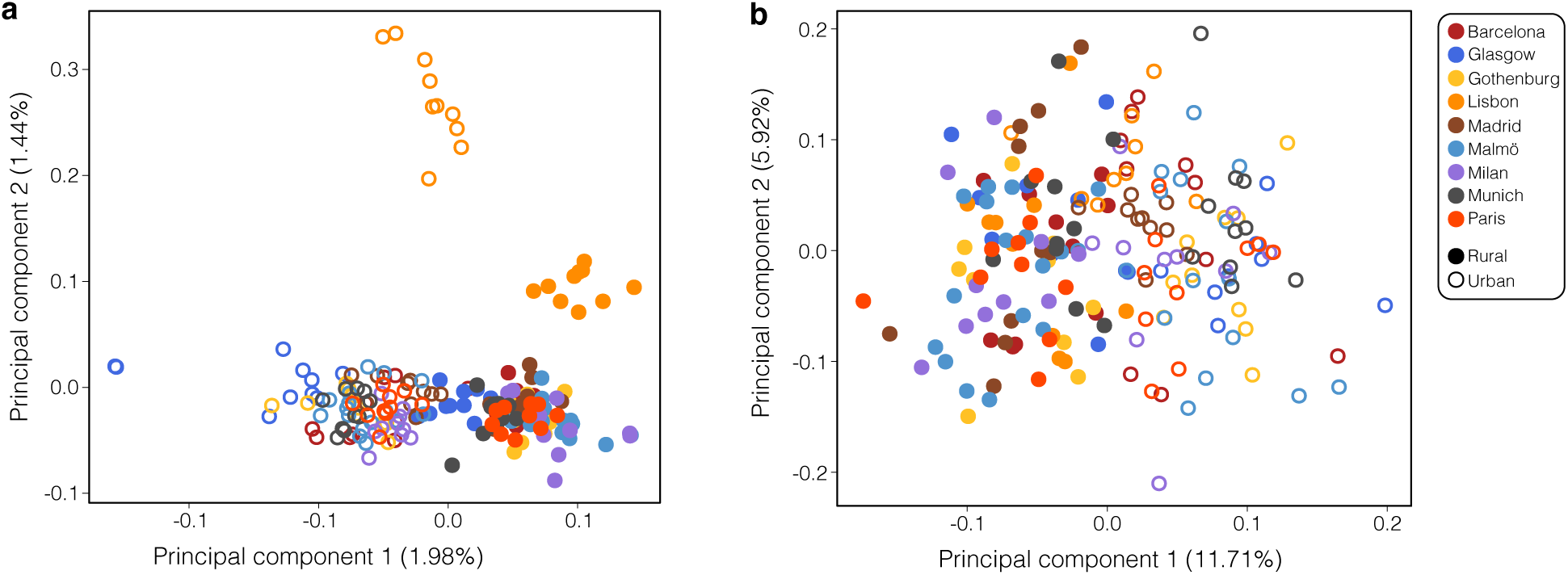
PCA based on *LFMM* and “*Core urbanisation SNPs*”. The first two principal component axis are shown (Percentage of total variance explained), based on a, *LFMM* candidate SNPs (N=2,758) and b, shared “*core urbanisation SNPs*” between *LFMM* and *BayPass* (N=34).

**Fig. S5.**
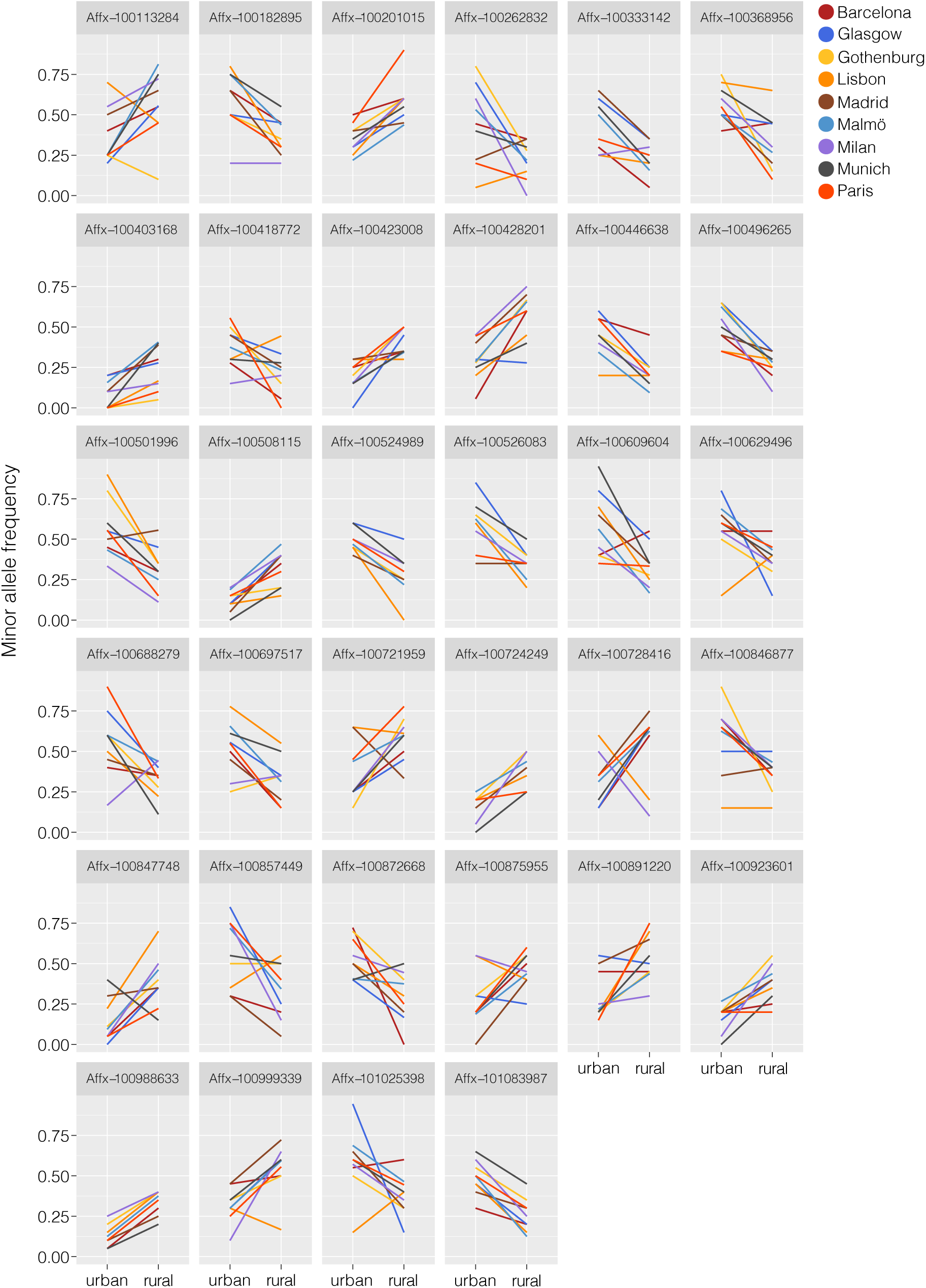
Differences in allele frequency between urban and rural populations in the “*core urbanisation SNPs*”. Minor allele frequency (MAF) trajectories between urban and rural populations for all “*core urbanisation SNPs*” (N = 34). The Affx-number in the header shows the name for each SNP on the Affymetrix SNP-chip used for the genotyping of all individuals in this study.

**Fig. S6.**
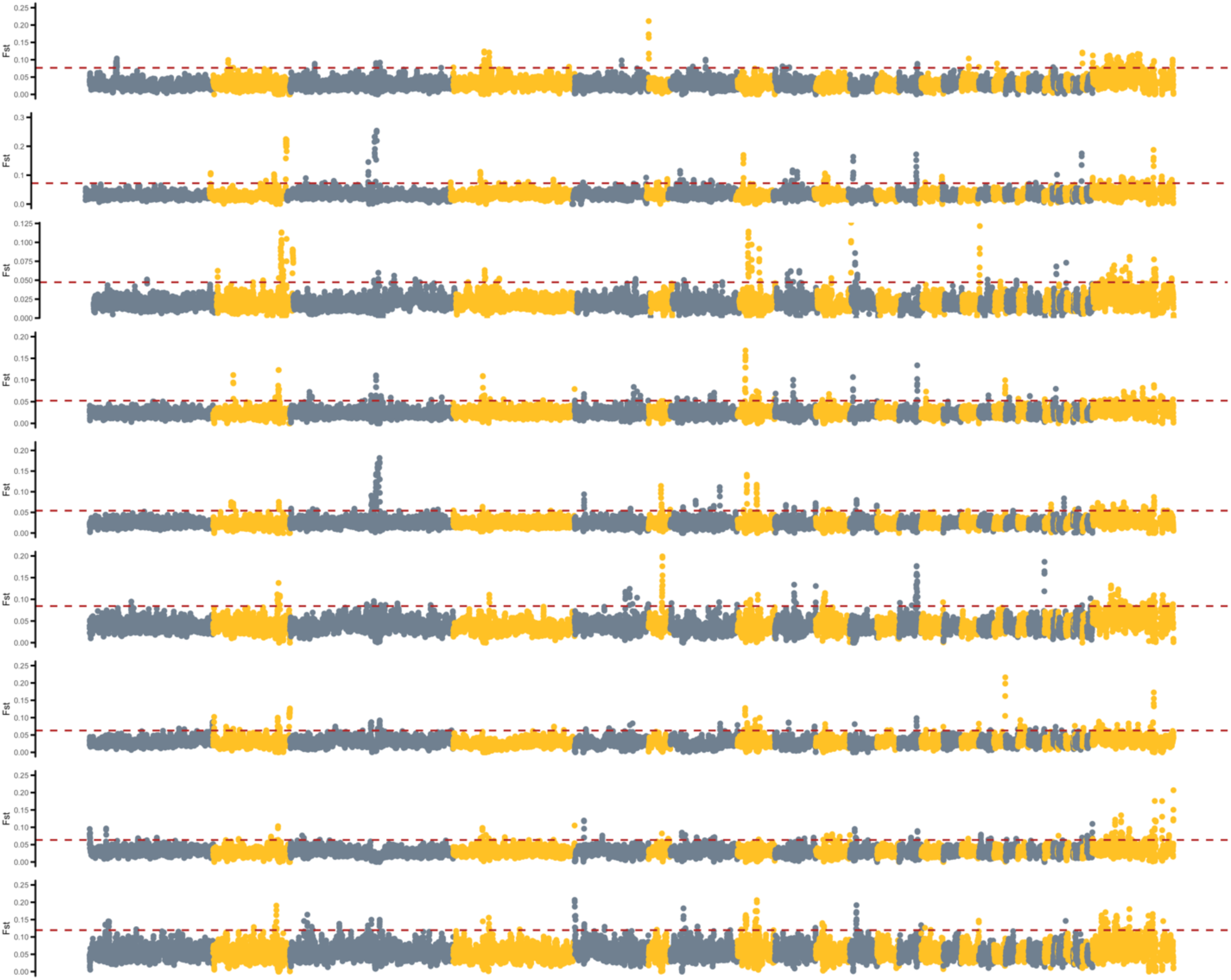
F_ST_ Manhattan plots. Manhattan plots showing the F_ST_ values (200 kb sliding windows with 50 kb steps) between urban and rural individuals across the genome for each population pair (urban-centre). The red dashed line shows the 99^th^ percentile of the F_ST_ distribution for each population pair.

**Fig. S7.**
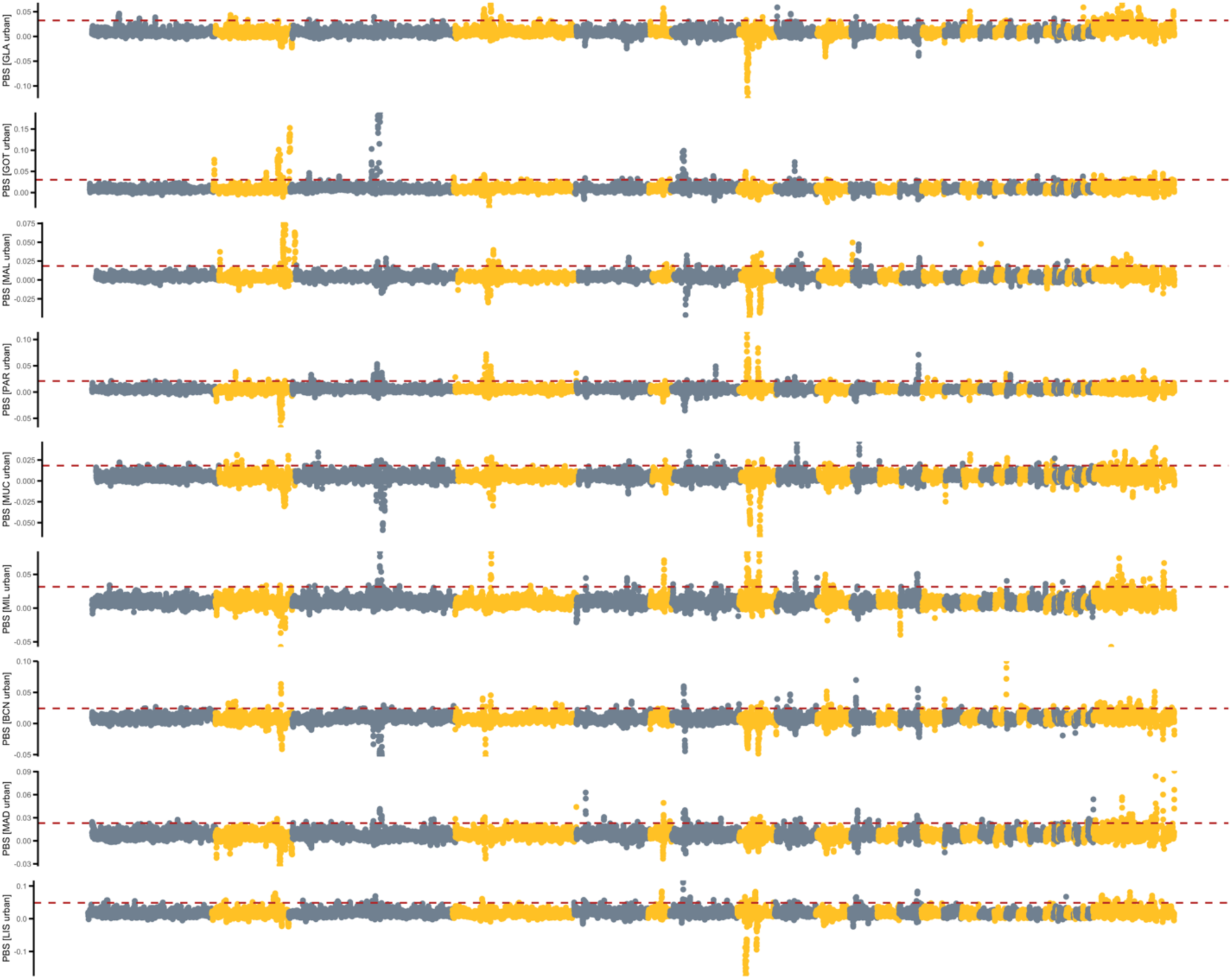
*PBS* Manhattan plots with LIS rural as outgroup. Manhattan plots showing the window-based *PBS* score (200 kb windows with 50 kb steps) between urban and rural individuals across the genome for each population pair (urban-centre). *PBS* scores were calculated using the rural population from Lisbon as an outgroup., *PBS* for Lisbon was calculated using GLA rural as the outgroup. The dashed lines show the 99^th^ percentile of the *PBS* distribution.

**Fig. S8.**
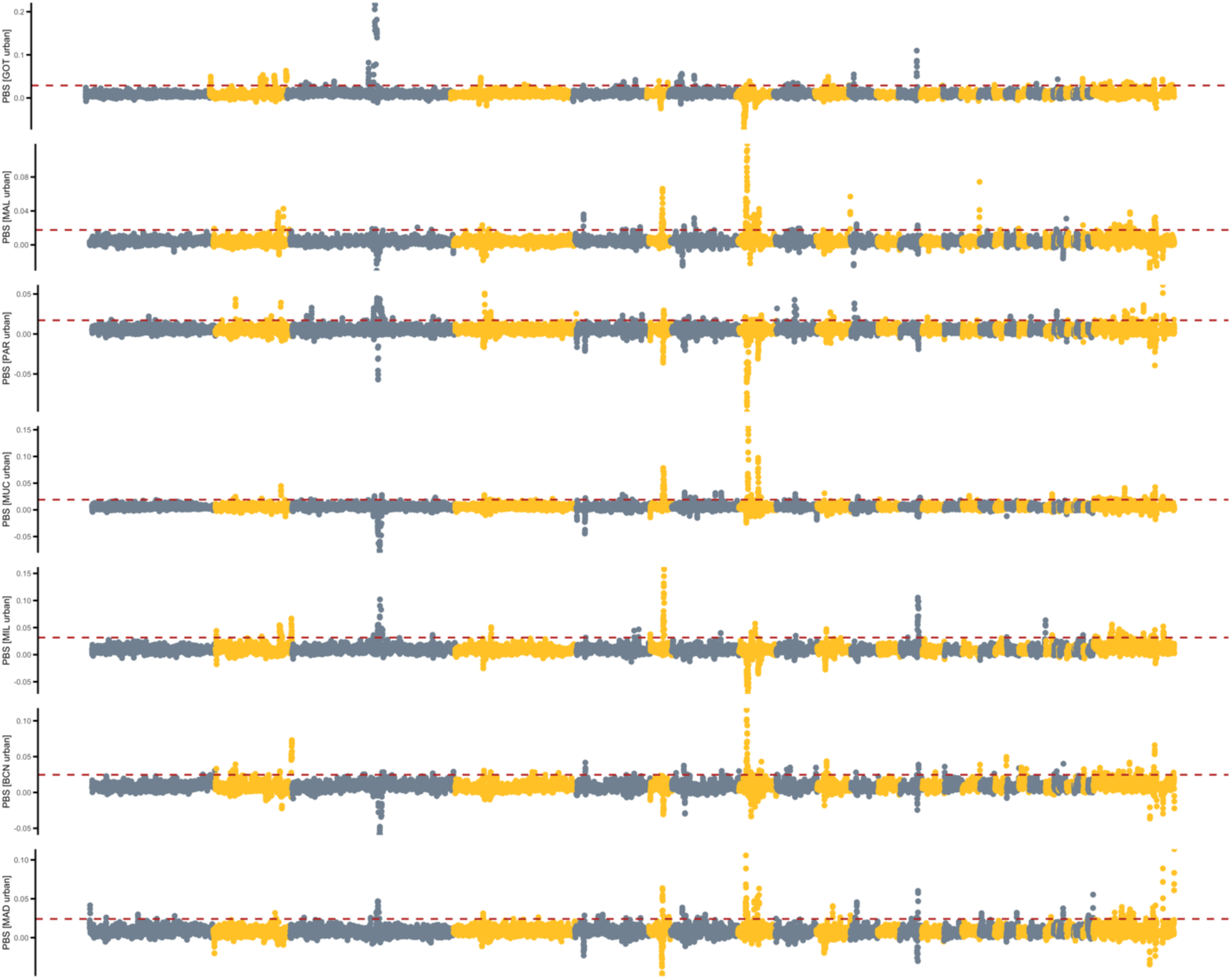
PBS Manhattan plots with GLA rural as outgroup. Manhattan plots showing the window-based *PBS* score (200 kb windows with 50 kb steps) between urban and rural individuals across the genome for each population pair (urban-centre). *PBS* scores were calculated using the rural population from Glasgow as an outgroup. PBS scores are shown for all population pairs except Glasgow and Lisbon. The dashed lines show the 99^th^ percentile of the PBS distribution.

**Fig. S9.**
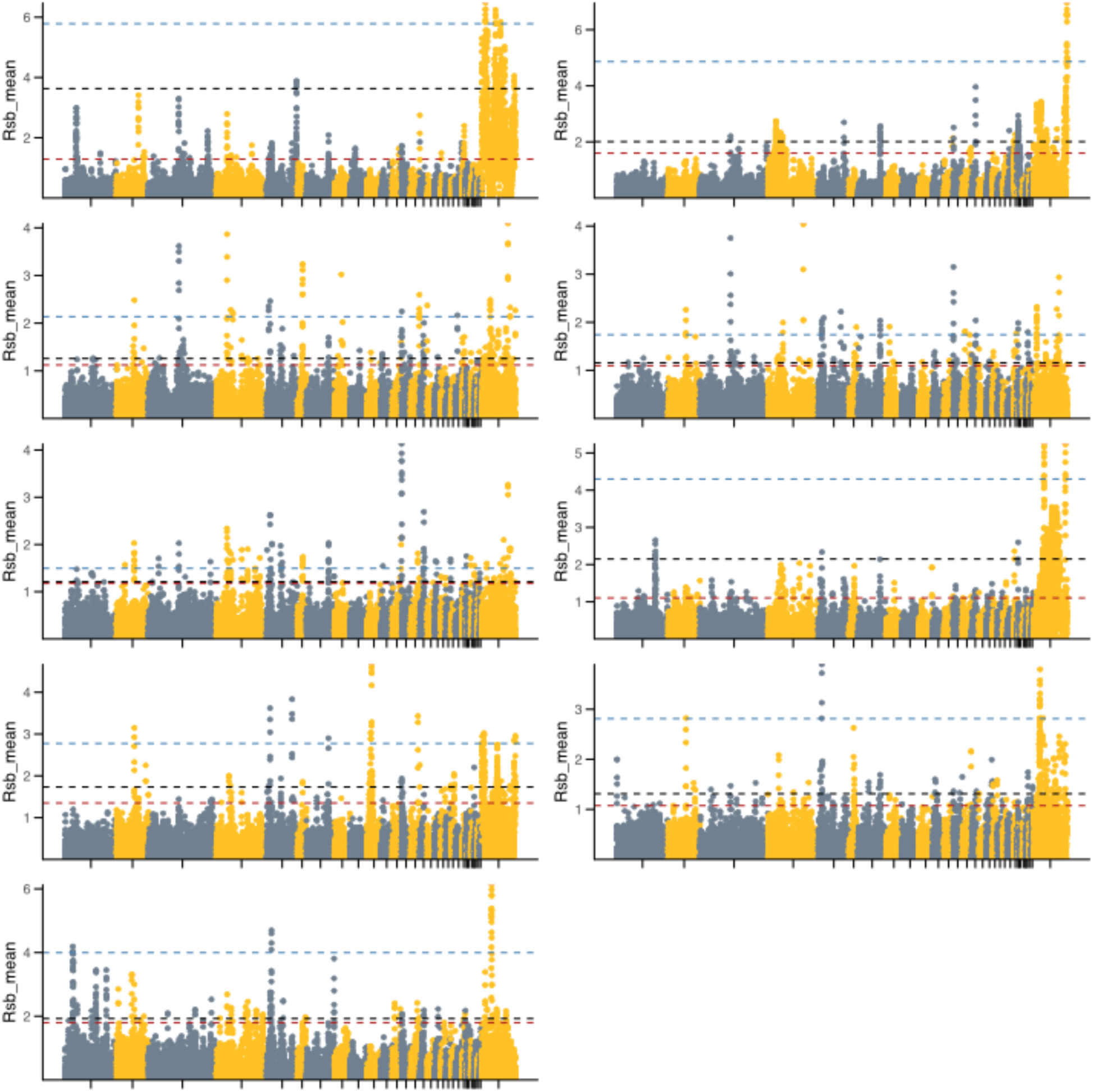
*Rsb* Manhattan plots. Manhattan plots showing the absolute haplotype-based selection scores (*Rsb* value in 200 kb windows with 50 kb steps) between urban and rural individuals across the genome for each population pair (urban-centre). The dashed lines show the 99^th^ percentile of the *Rsb* distribution for all chromosomes (black), only autosomes (red) and only the Z chromosome (blue). We selected outlier SNPs as those with SNP-based *Rsb* values above 4.

**Fig. S10.**
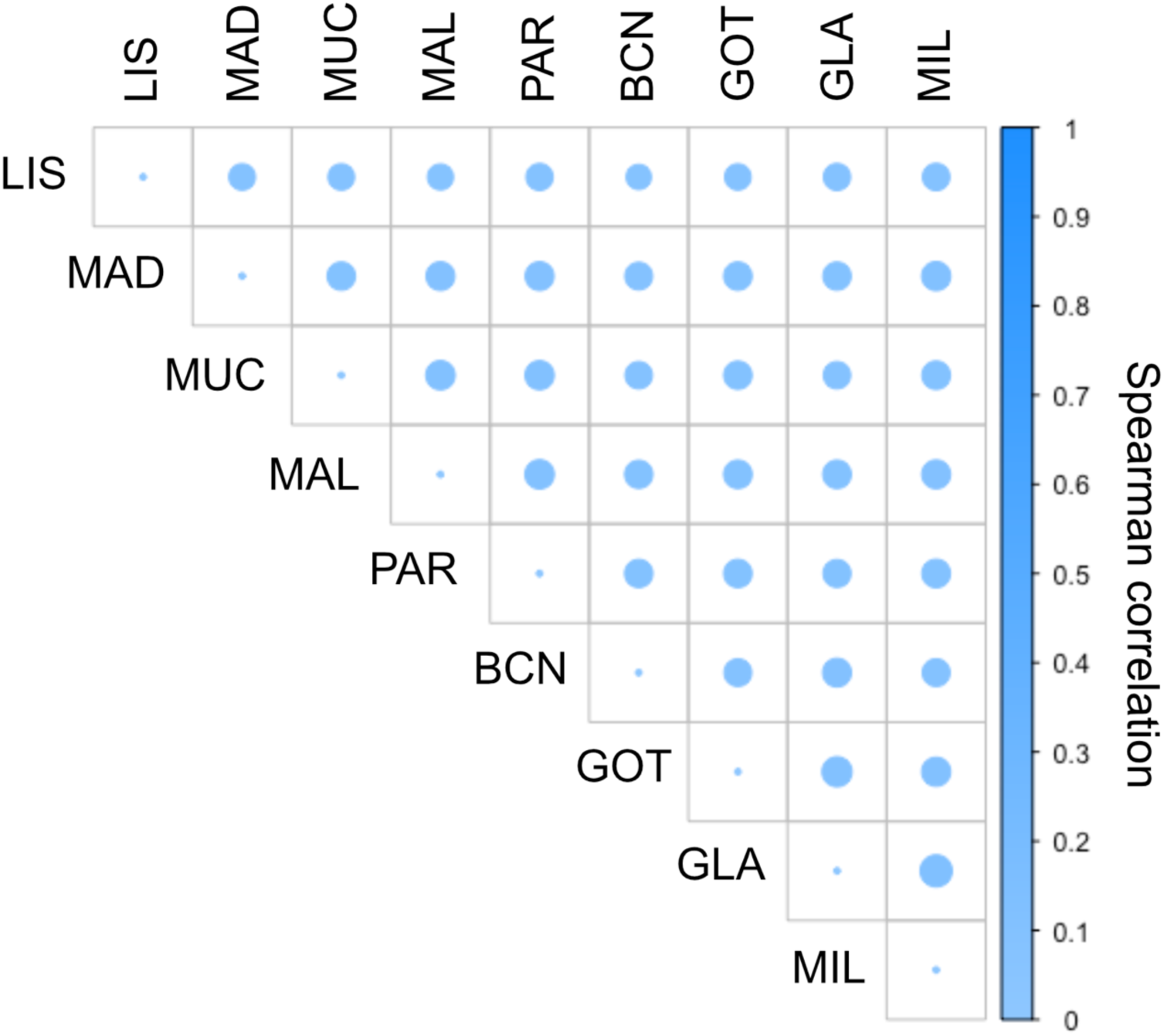
Correlation plot of absolute *Rsb* values across populations. Shown are spearman correlations between absolute SNP-based *Rsb* values across all population pairs. Spearman correlations were uniformly low across populations, ranging from 0.08 to 0.013.

**Fig. S11.**
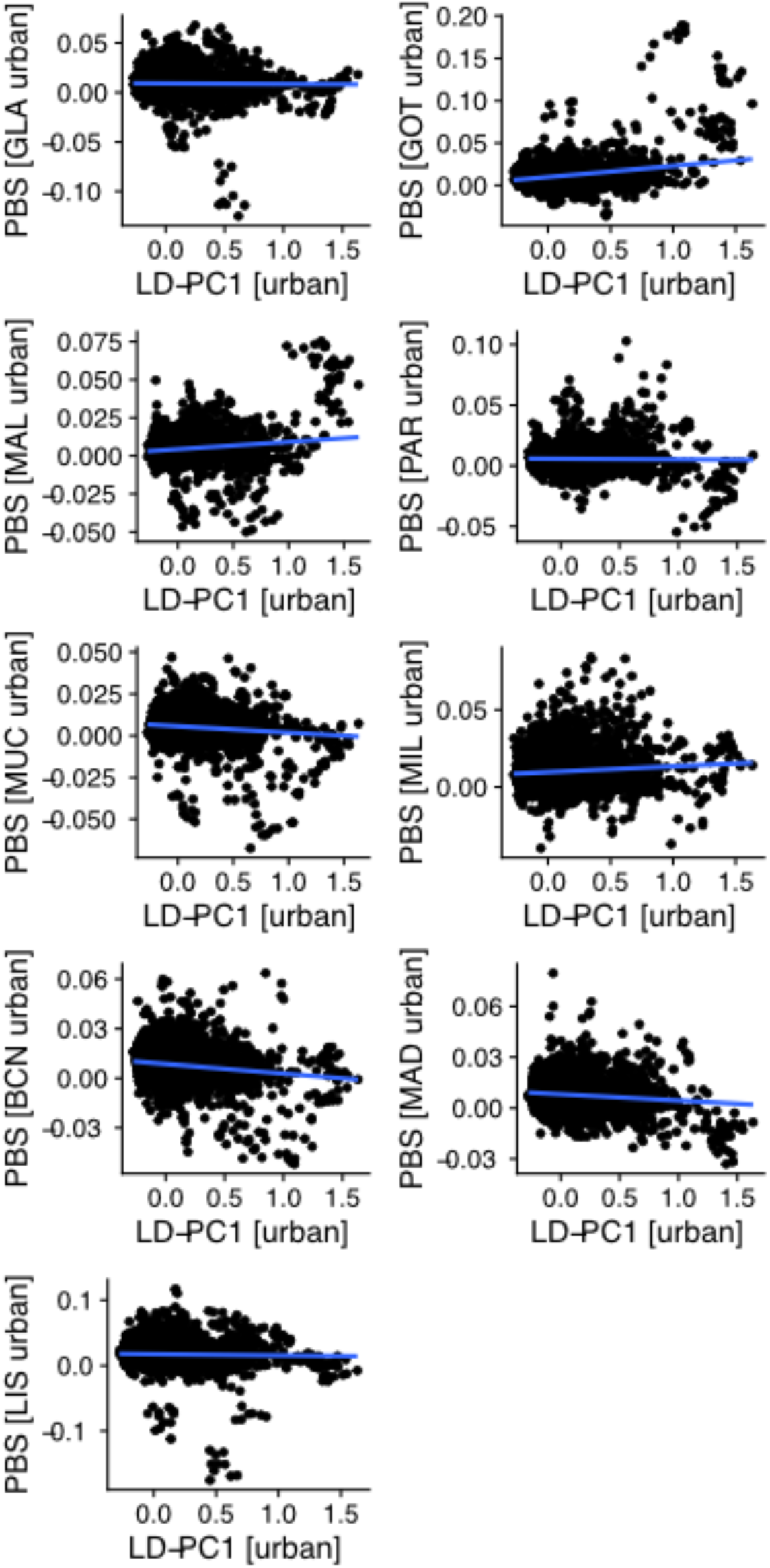
Correlation of *PBS* and *LD*_*urb*_*-PC1* for each population. LD was calculated and summarised across all urban populations.

**Fig. S12.**
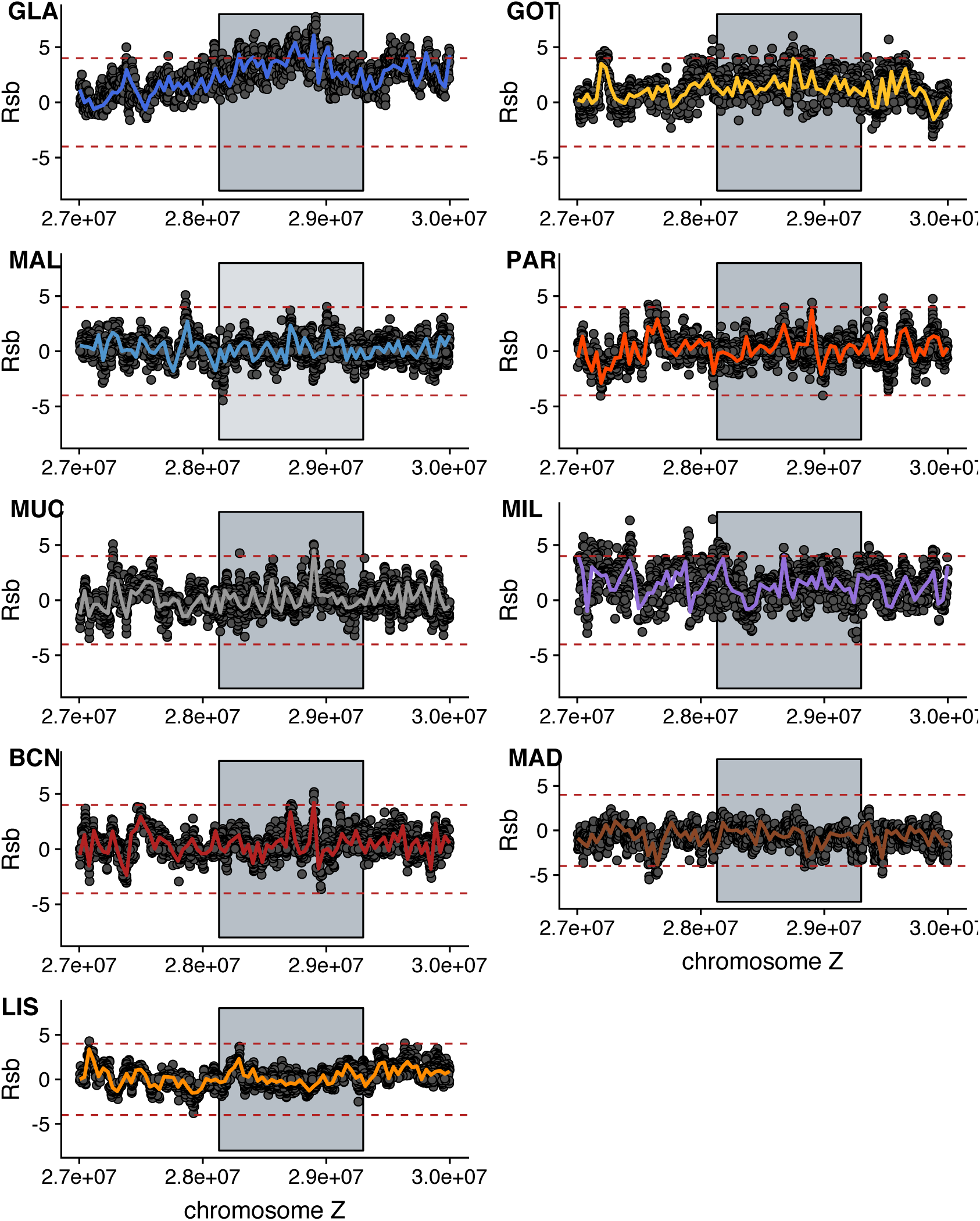
Signatures of selection around the *PTPRD* gene. *Rsb* scores for each SNP around and within the *PTPRD* gene (grey box) on the Z chromosome by locality, including loess smoothed values (span = 0.2). The upper and lower dashed lines show significance thresholds for signs of selection, respectively. BCN: Barcelona; GLA: Glasgow; GOT: Gothenburg; LIS: Lisbon; MAD: Madrid; MAL: Malmö; MIL: Milan; MUC: Munich; PAR: Paris.

**Fig. S13.**
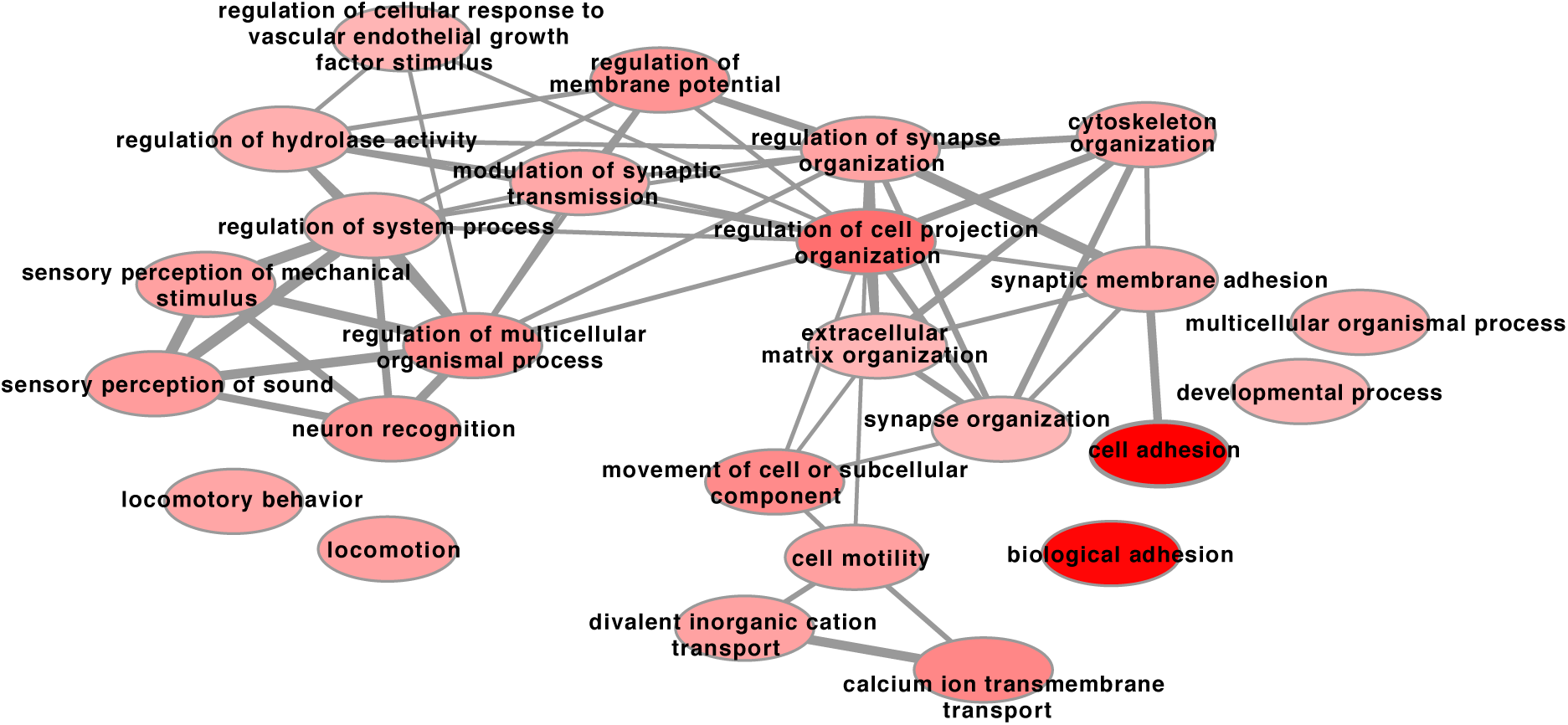
Network of urbanisation associated (*LFMM*) gene ontology (GO) terms (biological processes). The network shows the relationship of GO terms associated with urban-associated genes based on the number of shared genes between GO terms. The number of shared genes is given by the thickness of the grey lines. The colour intensity shows the degree of enrichment based on the p-value, with the highest intensity showing the lowest p-value. 28 GO terms were enriched at a false discovery rate (FDR) < 0.05 and 36 GO terms had an FDR > 0.05 after correction for multiple testing with FDR values ranging from 1.87 × 10^−7^ to 2.23 × 10^−1^.

**Table S1.** Locality (city name), year of sampling, season, centred geographical coordinates per site (urban/rural populations), urbanisation degree (PC_Urb_, positive values indicate higher urbanisation intensity), number of genotyped individuals (n), expected heterozygosity (H_e_), pairwise genetic differentiation (F_ST_) and distance between the urban and rural populations for each of the studied localities. Cities are sorted in alphabetical order.

| City | Year | Season | Urban | | | | Rural | | | | $F_{ST}$ | Distance (km) |
| --- | --- | --- | --- | --- | --- | --- | --- | --- | --- | --- | --- | --- |
| | | | Coordinates | $PC_{Urb}$ | $n$ | $H_e$ | Coordinates | $PC_{Urb}$ | $n$ | $H_e$ | | |
| Barcelona | 2015 | Winter | 41°23'24.0"N<br>2°11'24.0"E | 2.00 | 10 | 0.341 | 41°42'00.0"N<br>2°21'36.0"E | -2.29 | 10 | 0.348 | 0.012 | 5 |
| Glasgow | 2015 | Breeding | 55°52'48.0"N<br>4°15'36.0"W | 2.96 | 10 | 0.337 | 56°07'12.0"N<br>4°35'24.0"W | -1.71 | 10 | 0.344 | 0.013 | 33 |
| Gothenburg | 2015 | Breeding | 57°41'24.0"N<br>11°56'24.0"E | 2.13 | 10 | 0.342 | 57°30'00.0"N<br>12°00'36.0"E | -2.39 | 10 | 0.350 | 0.016 | 25 |
| Lisbon | 2014 | Post-Breeding | 38°44'24.0"N<br>9°10'48.0"W | 0.65 | 10 | 0.321 | 38°51'36.0"N<br>8°49'48.0"W | -1.86 | 10 | 0.333 | 0.050 | 33 |
| Madrid | 2014 | Breeding | 40°26'24.0"N<br>3°43'48.0"W | 1.60 | 10 | 0.344 | 40°34'12.0"N<br>4°09'36.0"W | -2.05 | 10 | 0.348 | 0.013 | 39 |
| Malmö | 2013 | Breeding | 55°36'00.0"N<br>12°59'24.0"E | 2.32 | 16 | 0.356 | 55°39'00.0"N<br>13°34'12.0"E | -2.44 | 16 | 0.355 | 0.009 | 37 |
| Milan | 2014 | Post-Breeding | 45°31'48.0"N<br>9°12'36.0"E | 1.49 | 10 | 0.342 | 45°49'12.0"N<br>9°17'24.0"E | -2.24 | 10 | 0.343 | 0.026 | 34 |
| Munich | 2015 | Winter | 48°07'48.0"N<br>1°34'12.0"E | 3.39 | 10 | 0.350 | 47°58'12.0"N<br>11°14'24.0"E | -1.36 | 10 | 0.351 | 0.004 | 30 |
| Paris | 2014 | Breeding | 48°52'12.0"N<br>2°10'48.0"E | 2.36 | 10 | 0.351 | 48°18'00.0"N<br>2°39'36.0"E | -2.58 | 10 | 0.351 | 0.004 | 70 |

Notes: Expected heterozygosity is significantly lower in urban compared to rural great tits in Glasgow, Gothenburg Barcelona, Madrid and Lisbon based on t-tests ( $P < 0.05$ ).

**Table S2.** Genes associated with “*core urbanisation SNPs*”.

| **Gene symbol** | **Gene name/ Description** | **Selection**<br>**(N. localities)** |
| --- | --- | --- |
| *FARP1* | FERM, ARH/RhoGEF and pleckstrin domain protein 1 | 1 |
| *CELF2* | CUGBP Elav-like family member 2 | 0 |
| *IQGAP2* | IQ motif containing GTPase activating protein 2 | 1 |
| *RNF38* | ring finger protein 38 | 2 |
| *DACH1* | dachshund family transcription factor 1 | 0 |
| *DHCR24* | 24-dehydrocholesterol reductase | 0 |
| *SV2C* | synaptic vesicle glycoprotein 2C | 1 |
| *AVL9* | AVL9 cell migration associated | 0 |
| *CADM2* | cell adhesion molecule 2 | 0 |
| *GABRG3* | gamma-aminobutyric acid type A receptor gamma3 subunit | 0 |
| *DNAI1* | dynein axonemal intermediate chain 1 | 2 |
| *NTRK2* | neurotrophic receptor tyrosine kinase 2 | 4 |
| *ABCB1* | ATP binding cassette subfamily B member 1 | 0 |
| *ELP6* | elongator acetyltransferase complex subunit 6 | 0 |
| *ADAMTS12* | ADAM metallopeptidase with thrombospondin type 1 motif 12 | 3 |

Notes: Selection (N.localities) – Number of urban populations (localities) a particular gene is under selection in based on the haplotype-based selection analysis (Rsb value)

**Table S3.**
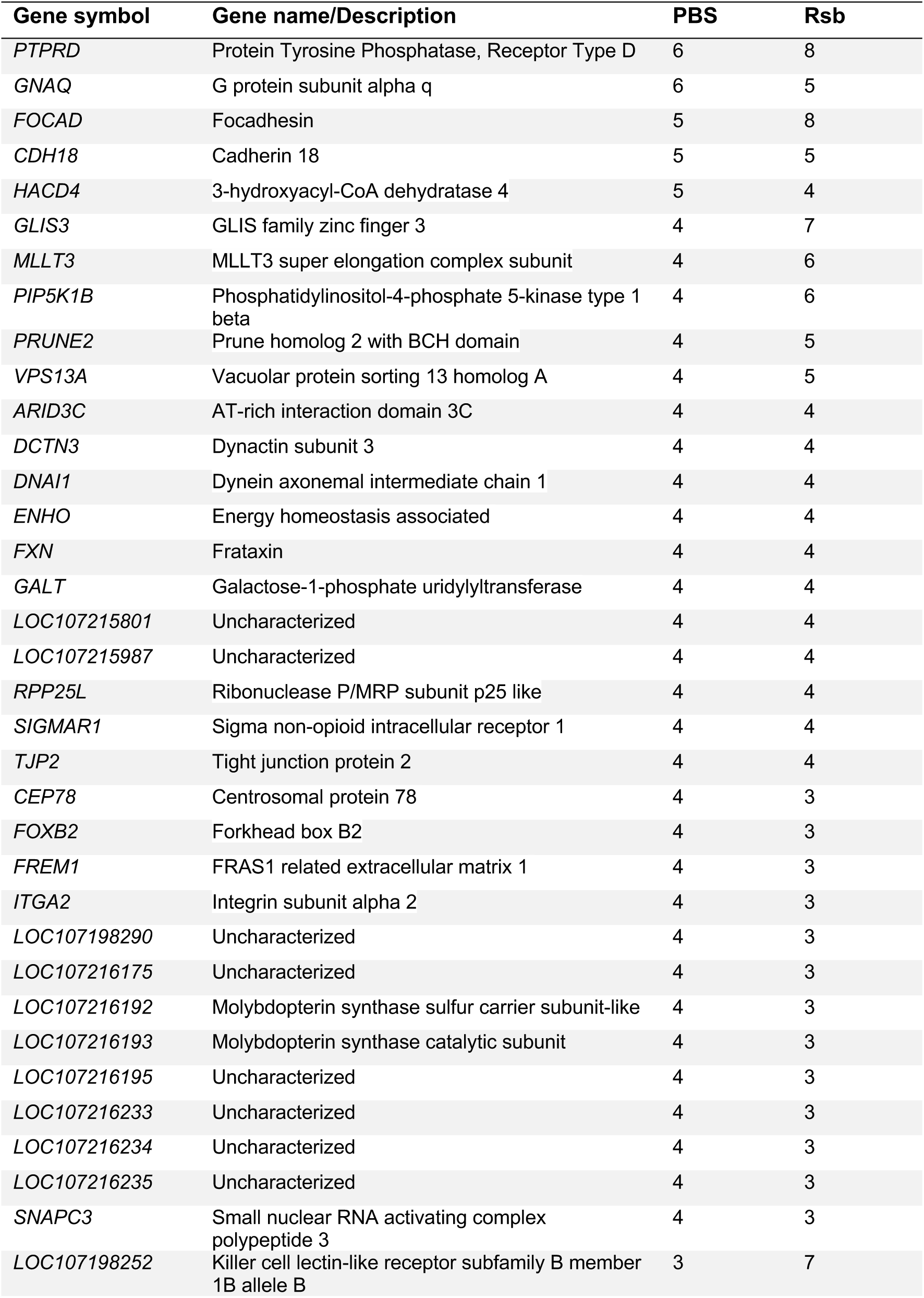
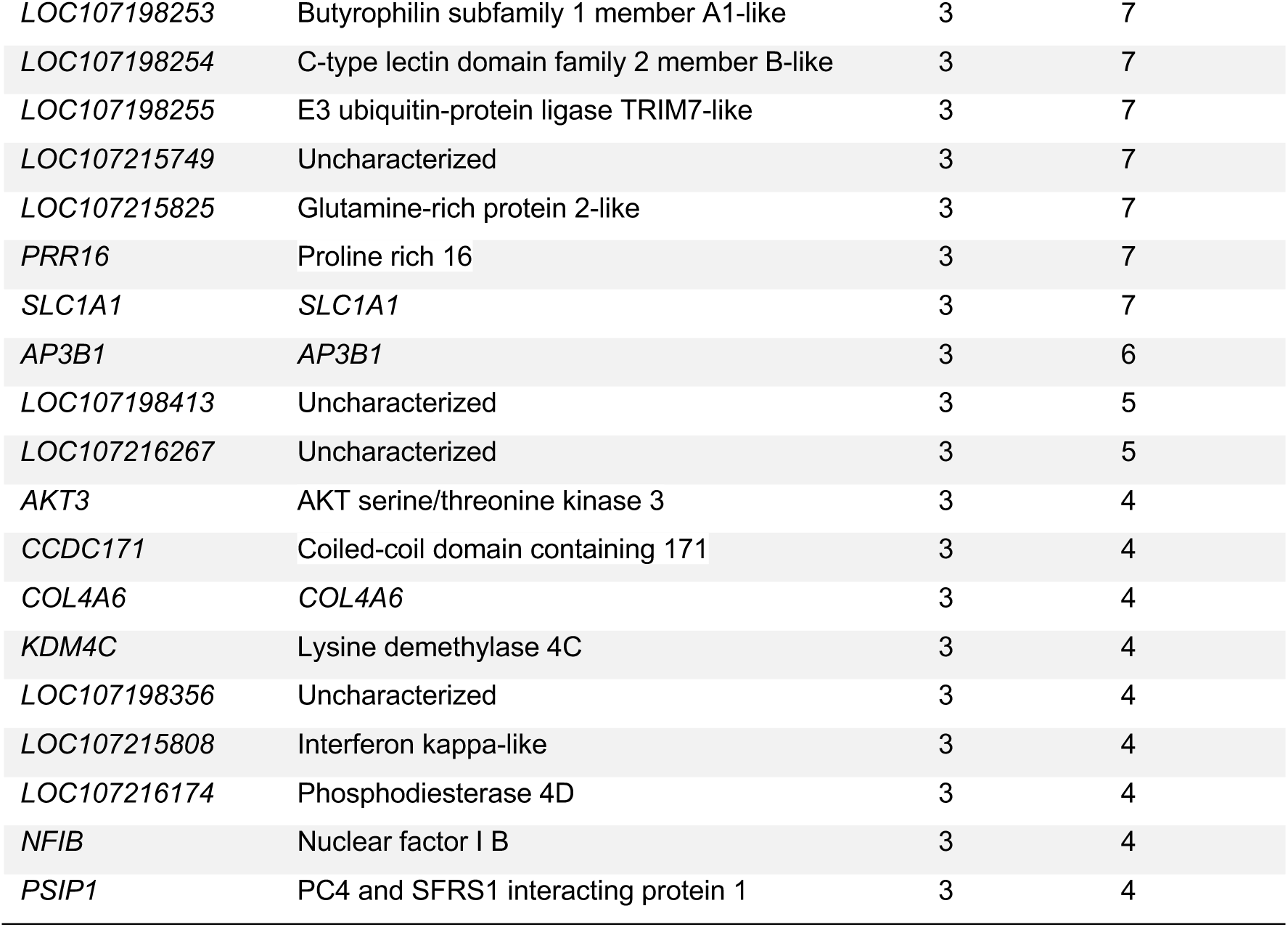
Genes putatively under selection in at least four cities based on *PBS* and/or *Rsb*. The number gives the number of cities in which a gene was detected to be under selection based on the given summary statistic.

| Gene symbol | Gene name/Description | PBS | Rsb |
| --- | --- | --- | --- |
| *PTPRD* | Protein Tyrosine Phosphatase, Receptor Type D | 6 | 8 |
| *GNAQ* | G protein subunit alpha q | 6 | 5 |
| *FOCAD* | Focadhesin | 5 | 8 |
| *CDH18* | Cadherin 18 | 5 | 5 |
| *HACD4* | 3-hydroxyacyl-CoA dehydratase 4 | 5 | 4 |
| *GLIS3* | GLIS family zinc finger 3 | 4 | 7 |
| *MLLT3* | MLLT3 super elongation complex subunit | 4 | 6 |
| *PIP5K1B* | Phosphatidylinositol-4-phosphate 5-kinase type 1 beta | 4 | 6 |
| *PRUNE2* | Prune homolog 2 with BCH domain | 4 | 5 |
| *VPS13A* | Vacuolar protein sorting 13 homolog A | 4 | 5 |
| *ARID3C* | AT-rich interaction domain 3C | 4 | 4 |
| *DCTN3* | Dynactin subunit 3 | 4 | 4 |
| *DNAI1* | Dynein axonemal intermediate chain 1 | 4 | 4 |
| *ENHO* | Energy homeostasis associated | 4 | 4 |
| *FXN* | Frataxin | 4 | 4 |
| *GALT* | Galactose-1-phosphate uridylyltransferase | 4 | 4 |
| *LOC107215801* | Uncharacterized | 4 | 4 |
| *LOC107215987* | Uncharacterized | 4 | 4 |
| *RPP25L* | Ribonuclease P/MRP subunit p25 like | 4 | 4 |
| *SIGMAR1* | Sigma non-opioid intracellular receptor 1 | 4 | 4 |
| *TJP2* | Tight junction protein 2 | 4 | 4 |
| *CEP78* | Centrosomal protein 78 | 4 | 3 |
| *FOXB2* | Forkhead box B2 | 4 | 3 |
| *FREM1* | FRAS1 related extracellular matrix 1 | 4 | 3 |
| *ITGA2* | Integrin subunit alpha 2 | 4 | 3 |
| *LOC107198290* | Uncharacterized | 4 | 3 |
| *LOC107216175* | Uncharacterized | 4 | 3 |
| *LOC107216192* | Molybdopterin synthase sulfur carrier subunit-like | 4 | 3 |
| *LOC107216193* | Molybdopterin synthase catalytic subunit | 4 | 3 |
| *LOC107216195* | Uncharacterized | 4 | 3 |
| *LOC107216233* | Uncharacterized | 4 | 3 |
| *LOC107216234* | Uncharacterized | 4 | 3 |
| *LOC107216235* | Uncharacterized | 4 | 3 |
| *SNAPC3* | Small nuclear RNA activating complex polypeptide 3 | 4 | 3 |
| *LOC107198252* | Killer cell lectin-like receptor subfamily B member 1B allele B | 3 | 7 |
| *LOC107198253* | Butyrophilin subfamily 1 member A1-like | 3 | 7 |
| *LOC107198254* | C-type lectin domain family 2 member B-like | 3 | 7 |
| *LOC107198255* | E3 ubiquitin-protein ligase TRIM7-like | 3 | 7 |
| *LOC107215749* | Uncharacterized | 3 | 7 |
| *LOC107215825* | Glutamine-rich protein 2-like | 3 | 7 |
| *PRR16* | Proline rich 16 | 3 | 7 |
| *SLC1A1* | *SLC1A1* | 3 | 7 |
| *AP3B1* | *AP3B1* | 3 | 6 |
| *LOC107198413* | Uncharacterized | 3 | 5 |
| *LOC107216267* | Uncharacterized | 3 | 5 |
| *AKT3* | AKT serine/threonine kinase 3 | 3 | 4 |
| *CCDC171* | Coiled-coil domain containing 171 | 3 | 4 |
| *COL4A6* | *COL4A6* | 3 | 4 |
| *KDM4C* | Lysine demethylase 4C | 3 | 4 |
| *LOC107198356* | Uncharacterized | 3 | 4 |
| *LOC107215808* | Interferon kappa-like | 3 | 4 |
| *LOC107216174* | Phosphodiesterase 4D | 3 | 4 |
| *NFIB* | Nuclear factor I B | 3 | 4 |
| *PSIP1* | PC4 and SFRS1 interacting protein 1 | 3 | 4 |

**Table S4.** Overrepresented GO terms for genes associated with urbanisation in the *LFMM* and *BayPass* analysis.

| **GO category** | **GO.ID** | **Description** | **Obs.** | **Fold enriched** | **p-value** | **FDR** |
| --- | --- | --- | --- | --- | --- | --- |
| **Processes** | GO:0098742 | cell-cell adhesion via plasma-membrane adhesion molecules | 38 | 2.920 | 8.27E-10 | 7.03E-07 |
| **Processes** | GO:0050808 | synapse organization | 55 | 2.078 | 1.04E-07 | 4.43E-05 |
| **Processes** | GO:0050803 | regulation of synapse structure or activity | 35 | 2.312 | 1.86E-06 | 4.36E-04 |
| **Processes** | GO:0016358 | dendrite development | 36 | 2.272 | 2.05E-06 | 4.36E-04 |
| **Processes** | GO:0042391 | regulation of membrane potential | 52 | 1.907 | 3.50E-06 | 5.95E-04 |
| **Processes** | GO:0050954 | sensory perception of mechanical stimulus | 24 | 2.399 | 4.06E-05 | 5.03E-03 |
| **Processes** | GO:0019932 | second messenger-mediated signalling | 46 | 1.823 | 4.14E-05 | 5.03E-03 |
| **Processes** | GO:0031346 | positive regulation of cell projection organization | 46 | 1.811 | 4.94E-05 | 5.25E-03 |
| **Processes** | GO:0010975 | regulation of neuron projection development | 56 | 1.691 | 5.63E-05 | 5.32E-03 |
| **Processes** | GO:0022604 | regulation of cell morphogenesis | 52 | 1.650 | 1.96E-04 | 1.63E-02 |
| **Functions** | GO:0045503 | dynein light chain binding | 8 | 4.440 | 1.98E-04 | 3.08E-02 |
| **Functions** | GO:0030507 | spectrin binding | 8 | 4.230 | 2.95E-04 | 3.08E-02 |
| **Functions** | GO:0050839 | cell adhesion molecule binding | 52 | 1.610 | 3.34E-04 | 3.08E-02 |
| **Functions** | GO:0005201 | extracellular matrix structural constituent | 23 | 2.090 | 4.86E-04 | 3.36E-02 |
| **Functions** | GO:0019199 | transmembrane receptor protein kinase activity | 15 | 2.520 | 6.22E-04 | 3.44E-02 |
| **Functions** | GO:0019838 | growth factor binding | 19 | 2.200 | 7.95E-04 | 3.67E-02 |
| **Functions** | GO:0005518 | collagen binding | 12 | 2.560 | 1.85E-03 | 7.33E-02 |
| **Functions** | GO:0051959 | dynein light intermediate chain binding | 7 | 3.530 | 2.40E-03 | 8.07E-02 |
| **Functions** | GO:0008066 | glutamate receptor activity | 7 | 3.380 | 3.18E-03 | 8.07E-02 |
| **Functions** | GO:0045505 | dynein intermediate chain binding | 7 | 3.380 | 3.18E-03 | 8.07E-02 |
| **Component** | GO:0097060 | synaptic membrane | 67 | 2.272 | 5.21E-11 | 8.86E-09 |
| **Component** | GO:0098984 | neuron to neuron synapse | 52 | 2.208 | 2.45E-08 | 2.08E-06 |
| **Component** | GO:0099572 | postsynaptic specialization | 48 | 2.061 | 7.54E-07 | 4.27E-05 |
| **Component** | GO:0098978 | glutamatergic synapse | 48 | 1.924 | 5.68E-06 | 2.00E-04 |
| **Component** | GO:0044309 | neuron spine | 28 | 2.450 | 5.89E-06 | 2.00E-04 |
| **Component** | GO:0033267 | axon part | 47 | 1.839 | 2.43E-05 | 6.89E-04 |
| **Component** | GO:0042383 | sarcolemma | 22 | 2.473 | 5.06E-05 | 1.23E-03 |
| **Component** | GO:0098793 | presynapse | 53 | 1.692 | 7.69E-05 | 1.64E-03 |
| **Component** | GO:0031594 | neuromuscular junction | 15 | 2.915 | 1.14E-04 | 2.16E-03 |
| **Component** | GO:0043235 | receptor complex | 43 | 1.712 | 2.87E-04 | 4.88E-03 |

Notes: Obs. (Total) – observed number of genes associated with gene ontology term.

